# Latent state selection, not outcome integration, resolves approach–avoidance conflict

**DOI:** 10.64898/2026.11.13.766373

**Authors:** Nathan J. Marchant, Nathan H.J. Pieterse, Gregorio Sonsini, Allison J. McDonald, Yvar van Mourik, Dustin Schetters, Taco J. De Vries

**Affiliations:** Amsterdam UMC location VUmc, Anatomy C Neurosciences, De Boelelaan 1108, 1081 HZ Amsterdam, The Netherlands; Amsterdam Neuroscience, Compulsivity Impulsivity and Attention, Amsterdam, The Netherlands; School of Pharmacy, Center for Neuroscience, University of Camerino, Via Madonna delle Carceri, 62032, Camerino MC, Italy

**Keywords:** <u>Key words</u>: anterior insula cortex, punishment, agency, choice, fiber photometry, hidden Markov models

## Abstract

Appropriate action selection under approach-avoidance is crucial for survival and emotional resilience. Tasks designed to measure this conflict typically observe response rates at an intermediate rate. However, whether this reflects a single compromised policy or alternation between opposing strategies is unresolved. We trained rats on a novel three trial operant conflict task and recorded anterior insular cortex (aIC) activity with fiber photometry. Fitting a cross-validated GLM-HMM to trial-by-trial choices identified three discrete latent states: a risk-taking state in which rats responded on most conflict trials, a risk-averse state in which they withheld on most conflict trials, and a perseverative state. Rather than integrating the two possible outcomes in conflict, aIC activity flexibly adjusted according to the active state, such that it resembled reward trials in the risk-taking state and punishment trials in the risk-averse state. We also found that acute alcohol increased punished responses by shifting state occupancy. In a yoked Pavlovian cohort, anticipatory aIC signals were reduced, indicating that this encoding requires instrumental agency. Together, these findings suggest that approach-avoidance conflict is resolved by selection between discrete strategies, and that insular recruitment of this process depends on the capacity to act.

**Significance statement:** Understanding the behavioural and neural mechanisms of decision-making in conflict is of importance from both theoretical and practical perspectives. Using a new operant conflict task in rats and a hidden Markov model of trial-by-trial choice, we show instead that rats alternate between discrete strategies that persist for tens of trials, approaching conflict in one and avoiding it in the other. Anterior insula activity followed the active strategy and required instrumental control. This provides a novel description of the computational mechanisms of conflict resolution, and indicates that anterior insula cortex is engaged by the strategy the animal is currently using, rather than by the conflict cue itself, and only when the animal can control the outcome.

## Introduction

Actions often carry the prospect of both reward and punishment, bringing the motivation to approach and to avoid into conflict. Resolving approach–avoidance conflict is a fundamental decision problem (Miller, 1944; Gray and McNaughton, 2000; McNally, 2021), and maladaptive conflict resolution is central to anxiety (Aupperle and Paulus, 2010; Pittig and Scherbaum, 2020) and substance use disorder (Deroche-Gamonet et al., 2004; Vanderschuren and Everitt, 2004; Jean-Richard-Dit-Bressel et al., 2018; Marchant et al., 2018).

Classic and value-based accounts propose that approach and avoidance tendencies sum into a single net value that determines action (Miller, 1944; Rangel et al., 2008; Levy and Glimcher, 2012; McNaughton et al., 2016). Consistent with this, net value signals have been described in ventromedial prefrontal and orbitofrontal cortex (Basten et al., 2010; Plassmann et al., 2010; Bartra et al., 2013) and in subcortical regions (Hulsman et al., 2024). Summation predicts graded compromise, with an intermediate response probability applied consistently across trials. However, at intermediate threat levels behaviour oscillates between the different options (Choi and Kim, 2010; Schlund et al., 2016; Verharen et al., 2019; Choi et al., 2022). These dynamics point to selection via mutual inhibition rather than summation (McNally, 2021).

Selection may also operate over longer timescales, as persistent strategies that determine how identical sensory inputs are translated into action (Flavell et al., 2022; Versteijnen and Heimel, 2026). Classical models of decision-making, including those recently applied to approach–avoidance decisions, assume a single, stationary strategy (Brown and Heathcote, 2008; Ratcliff and McKoon, 2008; Choi et al., 2022). However, more recent work using hidden Markov models, where each latent state has its own GLM (GLM-HMM), show that mice alternate between discrete strategies across many trials (Escola et al., 2011; Calhoun et al., 2019; Ashwood et al., 2022), and comparable state structure has been found across tasks and species (Calhoun et al., 2019; Marques et al., 2020; Bolkan et al., 2022; Ahmadlou et al., 2025), including rats, in which within-session shifts between reinforcement-learning strategies predicted response times and orbitofrontal encoding (Venditto et al., 2024).

The anterior insular cortex (aIC) is implicated in decisions that weigh reward against harm. It tracks risk (Kuhnen and Knutson, 2005; Preuschoff et al., 2008), is engaged during approach–avoidance decisions (Aupperle et al., 2015), is hyperactive in anxiety disorders (Paulus and Stein, 2006; Etkin and Wager, 2007), and in rodents supports risky choice and alcohol seeking despite punishment (Ishii et al., 2012; Seif et al., 2013; Campbell et al., 2019). As a salience-network hub proposed to mediate switching between brain states (Menon and Uddin, 2010), the aIC is positioned to track the prevailing strategy. In humans, a conflict cue evokes threat-like insula responses even when behaviour is reward-like, which has been attributed to salience coding (Chen et al., 2026). However, typically aIC signals are averaged across trials, and this cannot distinguish between integration, salience, or state selection three accounts.

A further question is whether aIC recruitment depends on the capacity to act. Insula activity is required to retrieve outcome values that guide goal-directed choice (Parkes and Balleine, 2013), and is associated with the sense of agency (Farrer and Frith, 2002) and the controllability of aversive events (Salomons et al., 2004). If aIC signals reflect the selection of an action strategy, they should therefore depend on instrumental control. Here, we recorded aIC activity with fiber photometry in rats performing an operant conflict task and fit a GLM-HMM to their trial-by-trial choices. Behaviour was best described by three latent states: Risk-taking, Risk-averse and Repeating. On conflict trials, aIC activity resembled reward trials in the Risk-taking state and punishment trials in the Risk-averse state. Acute alcohol increased punished responding by reducing occupancy of the Risk-averse state. In a yoked Pavlovian cohort, anticipatory and avoidance-related aIC signals were absent or reduced. Together, these results indicate that conflict is resolved by selection between discrete strategies, and that insular recruitment of this process depends on the capacity to act.

## Materials and Methods

### Subjects

We obtained 28 male Long-Evans rats, aged 8-10 weeks upon arrival (RjOrl:LE, RGD_151356971, Janvier Labs, France). All procedures were approved by the Vrije University Animal Welfare Body (Instantie voor Dierenwelzijn VU-VUmc (IvD)) and conducted under the authority of the Dutch Central Commission for Animal Research (CCD, permit #AVD1140020208565) in accordance with European law (Directive 2010/63/EU). Rats were divided into an Operant cohort (n=16) and a yoked Pavlovian cohort (n=12). After histological validation, 2 rats were removed from the Operant experiment and 3 from the Pavlovian experiment due to low jGCaMP7f expression or off-target fiber placement. This resulted in a final n=14 for the operant photometry analyses and n=9 for the Pavlovian photometry analyses. Behavioural analyses and the GLM-HMM included all rats that completed training (Operant n=16; Pavlovian n=12). One rat was removed from the operant cohort before the alcohol tests for health reasons, leaving n = 15 for the behavioural and latent-state analyses of the alcohol tests. This included the two rats excluded from the photometry analyses after histological validation. Of these, one rat completed five of the six test sessions (see Phase 4). The photometry analyses of the alcohol tests comprised n=13 rats. Only male rats were used because our food restriction protocol is designed for male rats.

### Housing conditions

Behavioural tests were conducted during the dark phase of the rat’s diurnal cycle (12h/12h). Upon arrival food and water were available ad libitum, however after recovery from surgery we started the food restriction protocol (see below). We housed the rats in groups of 2 or 3 throughout the experiment.

### Food Restriction and Weight Maintenance

To ensure consistent task motivation while strictly maintaining welfare, subjects were maintained on a rigorous, individualized food restriction protocol. Following a one-week acclimation period in our facility (on ad libitum chow), an individualized baseline free-feeding growth curve was established for each rat. Daily food rations were subsequently calculated based on age-adjusted metabolic energy requirements (e.g., 150 kcal/kg^0.75^ for growing animals, tapering to maintenance levels) to maintain subjects at approximately 85–90% of their projected free-feeding weight. To prevent age-related obesity, excessive fat deposition, and associated health complications, target body weights were statically capped once the animals reached 17 weeks of age.

Subjects were weighed a minimum of twice weekly. Daily caloric provisions were dynamically titrated based on individual metabolic demand, adherence to their specific growth curve, and daily task performance. Daily chow was administered in the home cage no earlier than 30 minutes following the completion of behavioural testing to maintain operant motivation. Although rats were group-housed, targeted individual feeding protocols were strictly implemented if cage mates exhibited disparate weight gain or deviated from their projected growth trajectories, ensuring precise caloric intake for every subject.

### Apparatus

We used Med Associates operant chambers for these experiments. Each operant chamber consisted of one wall with a retractable lever, and above the lever there were two cue lights, one white cue light and one red cue light. The other wall was equipped with a receptacle to deliver food pellets (Dustless precision pellets (F0021), 45 mg, Bio-Serve, USA), via an automated hopper connected to the receptacle by tubing. The grid floor with 19 rods (VFC-005) was connected to aversive stimulator/scrambler module to deliver footshocks (variable intensity, always 0.5 sec duration). All the protocols in the chambers were run by MedPC program, and the fan remained on for the whole duration.

For fiber photometry, excitation and emission light was relayed to and from the animal via optical fiber patch cord (0.48 NA, 400 µm flat tip; Doric Lenses). Blue excitation light (490 or 470nm LED [M490F2 or M470F2, Thorlabs]) was modulated at 211 Hz and passed through a 460-490nm filter (Doric Lenses), while isosbestic light (405nm LED [M405F1, Thorlabs]) was modulated at 531 Hz and passed through a filter cube (Doric Lenses). GCaMP7f fluorescence was passed through a 500-550nm emission filter (Doric Lenses) and onto a photoreceiver (Newport 2151). Light intensity at the tip of the fiber was measured before training sessions 2 times a week and kept at 40-50 µW. A real-time processor (RZ5P, Tucker Davis Technologies) controlled excitation lights, demodulated fluorescence signals and received timestamps of behavioural events. Data was saved at 1017.25Hz and analyzed with custom-made Matlab scripts.

### Viral vectors

We purchased premade viral vectors from the University of Zurich viral vector core: ssAAV-9/2-mCaMKIIα-jGCaMP7f-WPRE-bGHp(A) (jGCaMP7f). The titer injected was: jGCaMP7f, 4.0x10^12 gc/ml.

### Surgery

Both one day and thirty minutes prior to surgery we injected rats with the analgesic Carprofen (5 mg/kg, s.c.; Merial, Velserbroek, The Netherlands), we also administered Buprenorphine 30 minutes prior to surgery (0.01 mg/kg, s.c.). Surgery was performed under isoflurane gas anesthesia (PCH; Haarlem). We placed the anesthetized rat in a stereotactic frame (David Kopf Instruments, Tujunga, CA) and injected Lidocaine 10 mg/kg (Fresenius Kabi, Huis ter Heide, The Netherlands; RVG 51674) into the incision site prior to the incision. A craniotomy above aIC was performed, and 0.5 µl of AAV solution was injected unilaterally (hemisphere counterbalanced) into aIC (AP: +2.8, ML: +4.0 (2^◦^ angle), DV: -5.8 mm from Bregma) over 5 min. The needle was left in place for an additional 5 min. One 400μm optic fiber (Doric Lenses) was then implanted above aIC (AP: +2.8, ML: +3.8 (0^◦^ angle), DV: -5.5 mm from Bregma) and secured to the skull using dental cement (Antibiotic Simplex; Stryker, orthopaedics) and jewellers screws (Jeveka; M1x2 stainless steel screws). Carprofen (5 mg/kg; s.c.) was administered for 2 days after the surgery. Rats recovered for 10-14 days before food restriction began. Recordings started in Phase 1, 3 weeks after virus injection, to allow jGCaMP7f expression.

### Trial design

Figure 1B shows the trial design. Each trial begins with illumination of the respective cue light, which is above the lever. The cue remains on for 10 seconds prior to the insertion of the lever into the chamber. If the lever is pressed within 30 seconds, then the lever is retracted, the cue light is turned off, and the respective outcome (food pellet or shock) is delivered. There was no response-dependent stimulus signalling outcome delivery. If there was no lever press in 30 seconds then the lever is retracted, and the cue light is turned off. The inter-trial interval (ITI) started when the lever was pressed or retracted in omitted trials. ITI was on a variable interval (VI) 30 second schedule, ranging from 15 to 45 seconds.

**Figure 1.**
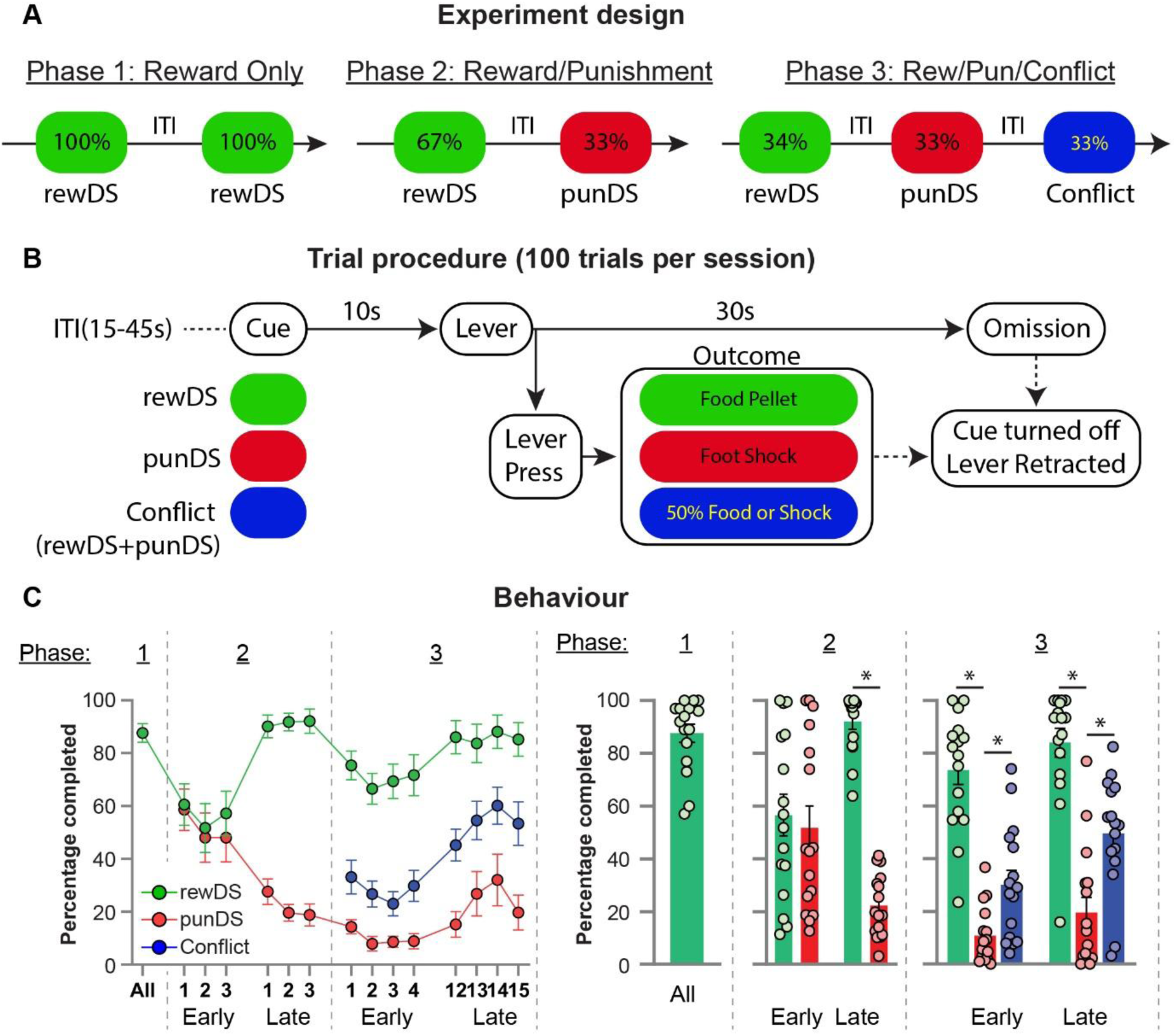
Task design and behaviour. **A.** Chronological progression of the experiment. Following recovery from AAV jGCaMP7f infusion and fiber implant, rats were trained through different phases of the operant discrimination task, in which 100 trials are given per session. In Phase 1 (3-8 sessions), there are only rewDS trials. In Phase 2 (6-15 sessions) 67% of trials are rewDS and 33% are punDS. In Phase 3 (15 sessions; 8 for one rat) there are three trial types: rewDS, punDS, and Conflict. **B.** Trial procedure. Each trial starts with a 10 second cue period, where the visual cue is illuminated but the lever is still retracted. After this, the lever is inserted for 30 seconds. If the rat presses the lever, the lever is retracted, the cue light is turned off, and the corresponding outcome is given (rewDS = food pellet; punDS = foot shock; Conflict = food pellet or foot shock). If the rat does not press the lever after 30 seconds, the lever is retracted, and the cue light is turned off. Following this is an inter-trial interval on a VI30 (15-45) second schedule. **C.** Mean (± SEM) response rate (%) across the three phases (n=16). Phases 2 and 3 are divided into ’early’ and ’late’ sessions. Phase 2: two-way RM ANOVA, Session × Cue interaction, F(1,15) = 132.8, p < 0.0001; rewDS vs punDS, early p = 0.24, late p < 0.0001. Phase 3: main effect of Cue, F(1.7,25.2) = 98.43, p < 0.0001; Session, F(1,15) = 4.40, p = 0.053; Session × Cue, F(1.6,24.7) = 1.50, p = 0.24. *, p < 0.05 (Šídák).

## Behavioural procedure

### Experiment 1 – Operant task

#### Experiment Design

Figure 1A shows the experiment design for the operant task. We trained rats 5 days a week, and a session lasted maximum 120 minutes, but typically 70 – 80 minutes depending on the number of trials completed versus omitted. We counterbalanced which cue (red or white) served as the reward discriminative stimulus (rewDS) or the punishment discriminative stimulus (punDS). Rats were tethered and recorded in every session.

*Phase 0:*

Initial lever press training. We first trained the rats in 30-minute sessions where the rewDS cue was on for the entire session, and the lever was inserted into the chamber throughout. If the rat pressed the lever (FR1), a food pellet was delivered, and the lever was retracted for 5 seconds. We sometimes baited the lever with crushed food pellets. We gave rats between 5 – 8 sessions in this phase before progressing to the next phase. Rats were tethered to the fiber optics for select sessions to systematically habituate them to the weight and movement of the patch cord.

*Phase 1:*

Reward training. During this phase we gave the rats 100 reward trials in a session. Rats received 3–8 sessions in this phase (median 7).

*Phase 2:*

Punishment training. During this phase, the first 2 trials were always reward trials. Subsequent trials were randomly drawn from an array designating 67% as reward and 33% as punishment. Each session consisted of 100 trials in total. To account for individual differences in pain tolerance, each rat began Phase 2 with a baseline foot-shock intensity of 0.20 mA. Shock intensity was titrated exclusively between sessions (never within a session) to establish stable behavioural discrimination, aiming for response rates of approximately 80% on rewDS trials and 20% on punDS trials. Rats received 6–15 Phase 2 sessions (median 10).

*Phase 3:*

Conflict training. During this phase, the first 2 trials were reward trials, after which trials were drawn from an array designating 1/3 as reward, 1/3 as punishment, and 1/3 as conflict. For conflict trials, the outcome (food pellet or foot-shock) was determined pseudo-randomly using a balanced 8-item array in MedPC (four 0s, four 1s) drawn without replacement, ensuring a strict 50/50 probability. Shock titration continued between daily sessions to aim for a response rate near 50% in conflict trials; for example, if a rat’s overall rewDS response rate dropped excessively, the shock intensity was decreased for the following session, whereas if punDS responding remained too high, the intensity was increased. All rats received 15 Phase 3 sessions except one, which received 8 because it took longer to reach the Phase 1 criterion. For this rat, sessions 1–4 were defined as early and sessions 5–8 as late. Across rats and sessions of Phases 2 and 3, shock intensity ranged from 0.20 to 0.39 mA.

*Phase 4:*

Alcohol tests. During this phase, the task conditions were identical to Phase 3. We tested the effect of acute alcohol injection on responding in the task. Ethanol (70% v/v) was diluted in sterile saline to 31.7% v/v (0.25 g/ml) and injected intraperitoneally at 2 ml/kg (0.5 g/kg) or 4 ml/kg (1.0 g/kg). Saline was injected at 4 ml/kg. We tested each dose (Saline, 0.5 g/kg, 1.0 g/kg) in a counterbalanced test order. We then repeated the test conditions again to acquire sufficient photometry data under each test condition. We injected the rats 15 minutes prior to the test session, after which they were placed into the chamber and tethered to the fiber optic and the session was started. Each rat was scheduled for 6 test sessions (each dose twice), and there was no fewer than three days between the test sessions to ensure there was no carry over effects of the prior test condition. One rat completed only five sessions, so its 0.5 g/kg data come from a single session. During the days in between we trained the rats in a normal Phase 3 training session.

#### Experiment 2 – Yoked Pavlovian task

To disambiguate the neural correlates of active decision-making from the passive sensory experience of task stimuli, a separate cohort of rats (n=12) was trained in a yoked Pavlovian task. We employed an offline, session-matched yoking procedure. Each Pavlovian rat was explicitly paired with a counterpart rat from Experiment 1. The complete chronological log from the Operant counterpart, including trial type (rewDS, punDS, Conflict), trial order (from trial 1 – 100), specific inter-trial intervals between each trial, exact response latencies, and trial outcomes (press vs. omission; reward vs. shock), was extracted and hard-coded into the relevant MedPC program for the paired Pavlovian subject and session. Thus, each rat in Experiment 2 progressed through the experimental phases experiencing the exact sequence and timing of sensory events generated by their Operant counterpart, but entirely independently of their own behaviour. In addition, shock intensity for each session was also set according to the shock intensity of the Operant counterpart (range 0.20 mA – 0.37 mA). The 12 operant partners were chosen to span the range of performance from the 16 operant rats.

*Session design:*

Because the Pavlovian cohort was trained for a total of 20 sessions, the distribution of training phases was proportionally scaled to preserve the relative experience of task acquisition and stabilization. The 20 Pavlovian sessions were divided into Phase 1 (3 sessions), Phase 2 (7 sessions), and Phase 3 (10 sessions). To map the Operant data onto this compressed 20-session Yoked schedule, we omitted the intermediate transitional days. Specifically, the 7 Yoked sessions for Phase 2 were derived directly from the first 3 (early) and last 4 (late) sessions of the Operant counterpart’s Phase 2 training. Similarly, the 10 Yoked sessions for Phase 3 were generated from the first 5 (early) and last 5 (late) sessions of the corresponding Operant Phase 3 log. This ensured the yoked rats were exposed to a representative balance of behavioural exploration and stabilized approach/avoidance outcomes.

*Trial design:*

Trials were identical to the operant paradigm. A trial commenced with a 10-s cue presentation, followed by lever insertion. The lever remained extended for a duration exactly matching the response latency of the paired Operant rat on that specific trial. If the Operant counterpart had responded, then for the Pavlovian Yoked rat the lever retracted, the cue turned off, and the corresponding outcome (food pellet or foot-shock) was delivered. If the Operant counterpart had omitted the trial, the lever retracted, and the cue turned off with no outcome delivered. For Pavlovian rats, "completed" and "non-completed" trials refer to the choice made by the yoked operant partner on that trial.

While instrumental lever presses by the Pavlovian rats were continuously monitored, they occurred at negligible rates. We measured magazine entries, quantified via infrared beam breaks in the food receptacle, as the primary behavioural metric to verify Pavlovian contingency learning. These were measured and are expressed as time (seconds) in the magazine over three 10 second epochs: Pre-Trial – prior to trial start; Cue – the 10 second cue period; Lever – 10 seconds after lever insertion. Because the response latency is inherently variable, the 10 Lever period contains many different events, therefore we also analysed magazine entries in the 10 seconds after outcome delivery. We always checked for leftover food pellets after each session and did not observe any.

### jGCaMP7f expression and fiber placement validation

We deeply anesthetized rats with isoflurane and Euthasol® injection (i.p.) and transcardially perfused them with ∼100 ml of normal saline followed by ∼400 ml of 4% paraformaldehyde in 0.1M sodium phosphate (pH 7.4). The brains were removed and post-fixed for 2 h, and then 30% sucrose in 0.1M PBS for 48 h at 4°C. Brains were then frozen on dry ice, and coronal sections were cut (40 µm) using a Leica Microsystems cryostat and stored in 30% sucrose in PBS stored at -20°C. We selected a 1-in-4 series and first rinsed free-floating sections (3 x 10 minutes) before incubation in PBS containing 0.5% Triton-X and 10% Normal Donkey Serum (NDS) and incubated for at least 48 h at 4°C in mouse anti-NeuN primary antibody (1:1000; Chemicon, MAB377). Sections were then repeatedly washed with PBS and incubated for 2-4 h in PBS + 0.5% Triton-X with 2% NDS and donkey anti-mouse secondary antibody DyLight 649 (1:500; Jackson ImmunoResearch, 715-495-150). After another series of washes in PBS, slices were stained with DAPI (0.1 ug/ml) for 10 min prior to mounting onto gelatin-coated glass slides, air-drying and cover-slipping with Mowiol and DABCO.

Slides were all imaged on a VectraPolaris slide scanner (VUmc imaging core) at 10x magnification, and QuPath was used for image analysis (Bankhead et al., 2017). Images containing aIC, from Bregma + 4.2 mm to +2.5 mm were identified and the boundary of expression for each rat was plotted onto the respective Paxinos and Watson atlas (Paxinos and Watson, 2008).

## Statistics

### Behaviour

Behavioural data and summary measures of photometry activity were analysed in GraphPad Prism (version 10.6.0 (890)). For all phases the dependent variable was the percentage of trials completed (i.e. lever pressed). Phases were analysed separately; Phase 1 was not analysed inferentially because only rewDS trials were presented. For Phase 2 we compared the first three (’early’) and last three (’late’) sessions of each rat; for Phase 3 we compared the first four (sessions 1–4) and last four (sessions 12–15) sessions. Data were analysed by two-way repeated-measures ANOVA with Session (early, late) and Cue (rewDS, punDS and, in Phase 3, conflict) as within-subject factors. Response rates by inferred latent state were analysed by two-way repeated-measures ANOVA with State (1–3) and Cue as within-subject factors. Where a design contained missing values (summary photometry measures for which a rat had fewer than five trials of a given type), a mixed-effects model fitted by restricted maximum likelihood (REML) was used instead. For within-subject factors with more than two levels, the Geisser–Greenhouse correction was applied. Significant main effects or interactions were followed by Šídák-corrected pairwise comparisons, with each row and each column of the design treated as a separate family. α = 0.05.

### Alcohol tests

Data from the two test sessions of each condition were combined, and the first 15 minutes of each session were excluded from the main analysis (see Results and Fig. S5). Percentage of trials completed was analysed by two-way repeated-measures ANOVA with Dose (saline, 0.5 g/kg, 1.0 g/kg) and Cue (rewDS, punDS, conflict) as within-subject factors, with the Geisser–Greenhouse correction, followed by Holm–Šídák-corrected comparisons between doses within each trial type. State occupancy (the percentage of a rat’s classified trials in each state at each dose) was analysed in the same way, with State (1–3) in place of Cue. Because the three occupancies sum to 100% within each rat and dose, the main effect of Dose is zero by construction; only the State × Dose interaction and the comparisons between doses within each state are reported. All alcohol analyses were performed in GraphPad Prism and included n=15 rats (n=13 for the photometry analyses). α = 0.05.

### Photometry

Recorded signals were first downsampled by a factor of 64, giving a final sampling rate of 15.89 Hz. The 405nm isosbestic signal was fit to the 470/490nm calcium-dependent signal using a first order polynomial regression. A normalized, motion-artefact-corrected ΔF/F was then calculated as follows: ΔF/F = (490nm signal − fitted 405nm signal)/fitted 405nm signal. The resulting ΔF/F was then detrended via a 90s moving average, and low-pass filtered at 3Hz. Traces were collected based on three categories of events: Cue period in responded trials, Cue and Lever period in omitted trials, and surrounding a lever press.

For analysis on responded trials, ΔF/F from 10s before trial start (cues on) to the lever insertion (i.e. 10 second cue period) were collated. For omitted trials, ΔF/F from 10s before trial start (cues on) to the 5 seconds after lever retraction (i.e. 40 second trial period) were collated. These traces were then baseline-corrected using the baseline time-period -10 seconds to -5 seconds prior to trial start. For analysis of lever presses, ΔF/F from 50 s before to 15 s after the press was collated. Each trace was z-scored against the same baseline as the cue-aligned trace of its trial, −10 to −5 s before cue onset. Anchoring the baseline to cue onset ensured that it was always taken during the inter-trial interval and was identical for cue-aligned and press-aligned traces from the same trial. Press-aligned traces are shown from 10 s before to 10 s after the press.

We converted the data of each trace into z-score by subtracting the mean baseline activity and dividing by the standard deviation. For each sub-analysis, we generated the mean per rat and then expressed the traces as the mean of the subject means. All subsequent analyses were conducted using the subject means.

We used three approaches to analyse the resulting traces: bootstrapped confidence intervals (bCI), permutation tests (PT), and summary statistics. The rationale for bCI and PT are described in detail here: (Jean-Richard-Dit-Bressel et al., 2020; Alonso-Lozares et al., 2024; Alonso-Lozares et al., 2025; van Mourik et al., 2026).

Bootstrapped confidence intervals were used to determine whether calcium activity was significantly different from baseline (ΔF/F = 0). A distribution of bootstrapped means was obtained by randomly sampling from traces with replacement (n traces for that response type; 5000 iterations). A 95% confidence interval was obtained from the 2.5th and 97.5th percentiles of the bootstrap distribution, which was then expanded by a factor of √(n/(n − 1)) to account for narrowness bias. Permutation tests were used to compare calcium activity between conditions. At each time point the observed difference between condition means was compared with a null distribution of 1,000 differences obtained by randomly reassigning condition labels. Comparisons in which every rat contributed to both conditions (for example, rewDS vs punDS within a phase) used paired permutations. Comparisons between partly overlapping or independent sets of rats used unpaired permutations of rat means. P values were two-tailed. α = 0.05 was Bonferroni-corrected for the number of comparisons made on the same traces.

For both bootstrap and permutation tests, only periods that were continuously significant for at least 0.5 s (≥ 8 consecutive samples at 15.89 Hz) were considered significant. Significant periods are shown as bars beneath each trace, and their start and end times are reported in Tables 1–4 and Supplementary Table S1 as the times of the first and last significant sample relative to the alignment event. Summary statistics were used for subject-based analyses of predefined epochs: the mean z-ΔF/F over the 10-s cue period for responded trials, and over the 30-s lever period for omitted trials. These subject means were analysed as described under Behaviour.

### Latent state model of conflict behaviour

Trial-by-trial lever-press decisions during Phase 3 were modelled with a Bernoulli Generalized Linear Model - Hidden Markov Model (GLM-HMM) (Ashwood et al., 2022; Bolkan et al., 2022). This approach allows us to detect distinct, temporally continuous latent behavioural states. Each state operates as an independent GLM, defined by a unique set of regression weights corresponding to task inputs. The model tracks transitions between these states over time, allowing the weights, and thus the rat’s behavioural strategy, to dynamically vary between set strategies. For a given state k, the probability of a lever press response at trial t (y_t_ = 1) is given by the logistic function:

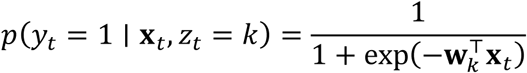

where y_t_ = 1 denotes a press, x_t_ is the input vector on trial t, z_t_ is the latent state and w_k_ is the weight vector of state k. States evolved as a first-order Markov chain with a stationary transition matrix, A_ij_ = p(z_t+1_ = j | z_t_ = i), and an initial-state distribution. Neither depended on trial inputs. The inputs were rewDS (1 on rewDS and conflict trials, otherwise 0), punDS (1 on punDS and conflict trials, otherwise 0), past pellet and past shock (1 if the previous trial in the same session ended in a pellet or a shock, otherwise 0; 0 on the first trial of a session), and a constant bias. Conflict trials were therefore represented as the conjunction of both cue inputs, with no separate conflict input. Weights are reported with positive values increasing the probability of a press. A single set of weights and transition probabilities was fitted to all rats. Weights had a weak zero-mean Gaussian prior (σ = 1,000, the ssm default); transitions had no prior. The model was fitted to all Phase 3 sessions (16 rats, 233 sessions, 23,126 trials). The number of free parameters was (K − 1) + K(K − 1) + 5K.

#### Model Fitting and Selection

Model performance was assessed by leave-one-rat-out cross-validation (16 folds). For each fold and each state range (1-5), models were refit on the training rats using an identical multi-start procedure. Held-out log-likelihood per trial was computed on the excluded rat by the forward algorithm. Choice prediction accuracy was computed one-step-ahead: the predicted state distribution at trial t was obtained by propagating the filtered posterior from trial t−1 through the transition matrix, so that predictions were conditioned only on prior choices and never on the choice being predicted. Additionally, the Akaike and Bayesian Information Criterion (AIC/BIC) were evaluated.

The 3-state GLM-HMM was compared against a single-state logistic GLM, with accuracy compared across rats by paired t-tests within each cue condition. To determine whether additional states reflected distinct strategies or subdivisions of existing ones, 4- and 5-state models were fit to the full dataset and evaluated in comparison to the 3-state model. We then compared the trial assignment and GLM weights using a posterior-probability assignment matrix (comparing 3 state solution to 4 state solution).

#### Trial State classification

The final three-state model was applied to all trials, and per-trial posterior state probabilities were computed with the forward–backward algorithm over each session (smoothed posteriors, p(z_t_ | y_1:T_)). For state-conditioned analyses, trials were assigned to the maximum-posterior state only when it’s probability was greater or equal to 0.65; lower-confidence trials were left unassigned. In a 3-state model (chance = 0.33), this threshold ensures that the assigned state is the decisively dominant latent strategy, while remaining lenient enough to account for the natural biological variance inherent in transitional behavioural periods.

For state-dependent photometry analyses, two rats that were included for the model fitting were excluded from subsequent analyses, leaving n=14. Subjects were required to have a minimum of 5 trials for a given cue-state combination to be included in the group-level analysis. Subjects failing to meet this threshold for a specific condition were excluded from that specific comparison to maintain a high signal-to-noise ratio in the averaged event-related transients. At this threshold, 11.8% of trials (2,731/23,126) were left unassigned.

#### State occupancy and dynamics

For each rat, occupancy was expressed as the percentage of classified trials assigned to each state. Within each session, classified trials were grouped into runs of the same state. A run could bridge gaps of up to three consecutive unclassified trials if the same state resumed on both sides. It ended when a different state was assigned, after more than three consecutive unclassified trials, or at the end of the session. Run length (dwell time) included bridged trials.

A state switch was counted between consecutive runs of different states, each at least three trials long; shorter runs were disregarded. To test how sensitive switch counts were to this criterion, the analysis was repeated with minimum run lengths of 1, 5 and 10 trials. A session was considered to contain a state if at least five of its trials were assigned to that state.

To estimate the timescale over which state allocation varied, a binary indicator of membership in each state (classified trials only) was decomposed into sums of squares between rats, between sessions within rats, and within sessions. Shock intensity was titrated only between sessions, so within-session variance cannot be attributed to changes in intensity.

Session occupancy (percentage of classified trials; sessions with at least 10 classified trials) was analysed with linear mixed models with a random intercept for rat, fitted by REML in statsmodels (Python). Change across Phase 3 was tested with occupancy ∼ session number. The relationship with shock intensity was tested with occupancy ∼ within-rat intensity + between-rat intensity + session number. Within-rat intensity was the session intensity minus the rat’s mean Phase 3 intensity; between-rat intensity was that mean. Intensity effects are expressed per 0.05 mA. As a robustness check, changes in occupancy between consecutive sessions were regressed on changes in intensity, again with a random intercept for rat.

#### Post hoc choice prediction from aIC activity

To test whether aIC activity carried choice information beyond the behavioural model, the fitted three-state GLM-HMM was held fixed (not refit) and used to generate a one-step-ahead prediction of the probability of responding on each trial, p^ₜ, by propagating the filtered state posterior from trial t−1 through the transition matrix. Predictions were therefore never conditioned on the choice being predicted; the smoothed posteriors used for state assignment elsewhere were not used for this analysis. Cue-period aIC activity was quantified on each trial as the mean z-ΔF/F over the 10 s cue period, and standardised within session. The log-odds of p^ₜ were entered as a fixed offset (coefficient fixed at 1) in a second-stage logistic regression to which aIC activity was added, such that aIC could explain only variance in choice not captured by the behavioural model. In its full form, the model was:

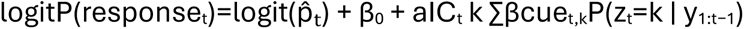

P(zₜ = k | y₁_:_ₜ₋₁) is the one-step-ahead probability of state k. To test whether aIC improved prediction overall, a model with a single aIC slope was compared with the offset-only model by leave-one-rat-out cross-validation (fitting on 13 rats and evaluating held-out log-likelihood per trial, using natural logarithms, on the 14th). The significance of this improvement was assessed by permutation (1,000 iterations), shuffling aIC values within rat × session × cue, which preserves cue-related differences in aIC while removing trial-level information. A model with separate aIC slopes for each cue type was evaluated by the same cross-validation, with improvement across rats tested by paired t-test. As a specificity control, these analyses were repeated using pre-trial activity (the 5 s preceding trial onset). Cue- and state-specific coefficients were estimated on the full dataset, with standard errors clustered by rat. Planned Wald contrasts compared conflict-trial slopes between the Risk-taking and Risk-averse states, and against the rewDS and punDS slopes within each state. The between-state contrast was additionally tested against a null distribution generated by shuffling state probabilities among conflict trials within session (1,000 iterations). Permutation p-values were computed as (number of permuted values ≥ observed + 1)/(number of permutations + 1), one-sided. Analyses were performed in Python using statsmodels and custom code.

#### Alcohol-test data and state decoding

The three-state GLM-HMM fitted to Phase 3 was applied to each alcohol-test session without refitting, with inputs coded as in Phase 3. Smoothed state probabilities were computed over the whole session, and a trial was assigned to its most probable state if that probability was at least 0.65 (11.7% of trials were unassigned). Sessions were then divided by trial start time into the first 15 min (≤ 900 s) and the remainder. State occupancy was the percentage of a rat’s assigned trials in each state, pooling its two sessions at each dose.

## Data and Code Availability

Trial-level behavioural data, subject-level photometry traces and model outputs, and Python code, including the GLM-HMM pipeline, is available at https://github.com/njmarchant/GLM-HMM_Conflict. MATLAB code for the photometry analysis are available at https://github.com/njmarchant/Photom-Anterior_Insula_Conflict.

## Results

### Acquisition of the Approach-Avoidance Conflict Task

To investigate the behavioural and neural dynamics of decision-making under approach-avoidance conflict, rats were trained on a multi-phase trial-based operant discrimination task (**Fig. 1A, B**). During the initial reward training (Phase 1), only rewarded trials were presented (rewDS), and rats rapidly acquired the instrumental lever press to obtain a food pellet (**Fig. 1C**). Following the introduction of randomly interleaved punishment trials (punDS) in Phase 2, rats initially failed to discriminate, responding equivalently on rewDS and punDS trials in early sessions (56.6% vs 51.7%; p = 0.24). By late Phase 2, responding was near-maximal on rewDS trials and suppressed on punDS trials (92.0% vs 22.3%; p < 0.0001; Session × Cue interaction: F(1,15) = 132.8, p < 0.0001; Fig. 1C). To establish approach–avoidance conflict (Phase 3), a third trial type was introduced (conflict), consisting of simultaneous presentation of both cues, which predicted a 50% probability of either outcome after a lever press. Shock intensity was titrated per rat between sessions to bring conflict responding towards 50%.

Responding differed between trial types throughout Phase 3, with conflict responding intermediate between rewDS and punDS responding in both early and late sessions (all pairwise comparisons p ≤ 0.0023; **Fig. 1C**). Conflict responding rose from 30.2% in early to 49.5% in late sessions, whereas rewDS (73.6% to 84.0%) and punDS responding (10.8% to 19.6%) changed less, although neither the main effect of Session nor the Session × Cue interaction reached significance (ps > 0.05).

### Anterior insular cortex activity emerges with discrimination learning and accompanies both responding and withholding

Throughout the task, population calcium activity was recorded from anterior insular cortex (aIC) using fiber photometry (**Fig. S1**). Time-locking aIC activity to trial onset in responded trials (**Fig. 2A**) revealed no significant deviation from baseline during Phase 1 (**Fig. 2B**). In contrast, following discrimination learning in Phase 2 (Phase 2 Late), trial start elicited sustained calcium transients on responded rewDS trials and punDS trials (**Fig. 2C**; **Table 1**). Mean cue-period activity increased from early to late Phase 2 (main effect of Session: F(1,13) = 26.15, p = 0.0002) and was higher on rewDS than punDS trials (main effect of Cue: F(1,13) = 8.68, p = 0.011; Session × Cue: F(1,13) = 1.21, p = 0.29; **Fig. 2D**). The permutation test did not identify a time window in which the rewDS and punDS traces differed (**Fig. 2C**), but mean cue-period activity was higher on rewDS trials in late (p = 0.0064) but not early Phase 2 (p = 0.12; **Fig. 2D**).

**Figure 2.**
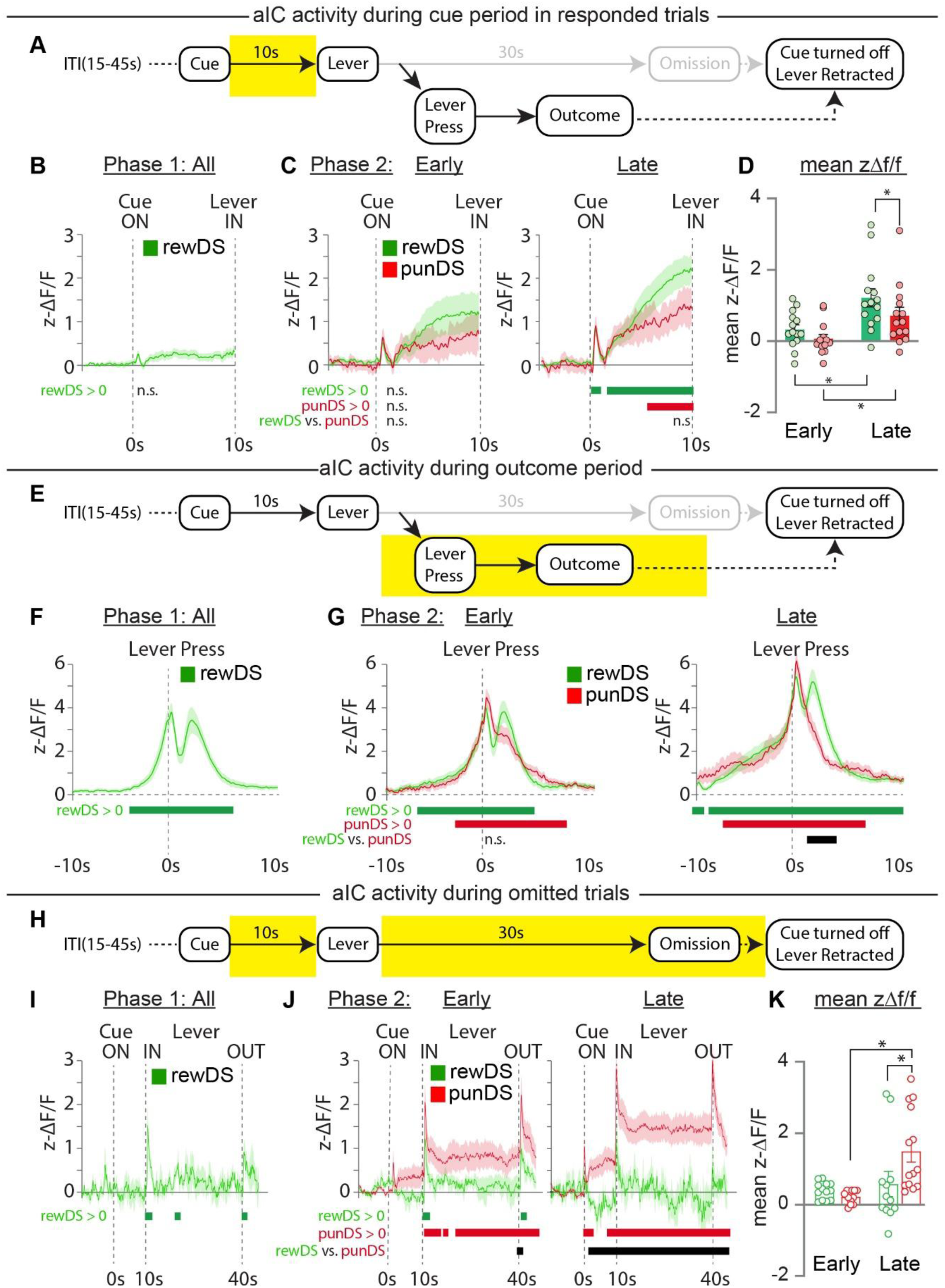
Anterior insular cortex activity emerges with discrimination learning and accompanies both responding and withholding. **A.** Trial procedure with the epoch of interest, the cue period, highlighted in yellow. B,C. Traces depicting aIC activity during the cue period (cue on = 0s) in responded trials. D Mean (± SEM) z-ΔF/F during the 10-s cue period. Activity increased from early to late Phase 2 (two-way RM ANOVA, main effect of Session: F(1,13) = 26.15, p = 0.0002) and was higher on rewDS than punDS trials (main effect of Cue: F(1,13) = 8.68, p = 0.011; Session × Cue: F(1,13) = 1.21, p = 0.29; rewDS vs punDS in late Phase 2: p = 0.0064). **E.** Trial procedure with the epoch of interest, the lever press, highlighted in yellow. **F, G.** Traces depicting aIC activity surrounding choice response (lever press = 0s). **H.** Trial procedure with the epoch of interest, the entire trial period in omitted trials, highlighted in yellow. **I,J.** Traces depicting aIC activity during the entire trial period (cue on = 0s) in omitted trials. **K.** Mean (± SEM) z-ΔF/F during the 30-s lever period of omitted trials (mixed-effects model: Session × Cue, F(1,8) = 5.41, p = 0.049; Session, F(1,13) = 9.86, p = 0.0078; Cue, F(1,13) = 2.35, p = 0.15; punDS vs rewDS in late Phase 2: p = 0.013; punDS, early vs late: p = 0.0008). For all trace graphs, the bars underneath represent the time along the trace where the statistical tests (bootstrapped CI, permutation tests) were significant. *Significant time windows are reported in Table 1. *, p < 0.05. n=14 rats, except K (rats with ≥ 5 omitted trials of a given type; early: rewDS n=13, punDS n=12; late: rewDS n=12, punDS n=14)*.

Aligning aIC activity to the execution of the decision (**Fig. 2E**) yielded a distinct biphasic calcium response for rewarded lever presses. This consisted of an initial transient corresponding to the action, followed by a second peak likely corresponding to reward retrieval (**Fig. 2F**). In late Phase 2, activity around rewDS and punDS presses was indistinguishable until after the press, suggesting a shared signature of decision commitment regardless of valence. The traces then diverged, coinciding with the second, reward-related peak (**Fig. 2G**; **Table 1**).

To isolate activity associated with withholding a response, aIC activity was examined during omitted trials (**Fig. 2H**). In Phase 1, omitted rewDS trials produced only brief transients at lever insertion and retraction (**Fig. 2I**; **Table 1**). In Phase 2, omitted punDS trials were already accompanied by activity above baseline from lever insertion onwards in early sessions (**Fig. 2J**). This activity increased with training (punDS, early vs late: p = 0.0008), and by late Phase 2 it exceeded activity on omitted rewDS trials after cue onset to the end of the trial, with only brief interruptions (**Fig. 2J**; **Table 1**). Mean lever-period activity showed a Session × Cue interaction (mixed-effects model: F(1,8) = 5.41, p = 0.049), with higher activity on omitted punDS than rewDS trials in late (p = 0.013) but not early Phase 2 (p = 0.65; **Fig. 2K**).

These data show that aIC is robustly recruited during decision-making phases of the trial as cognitive demand increases (i.e. Phase 2 discrimination after learning). Furthermore, this was true both for approach (responded rewDS trials) and avoidance choices (omitted punDS trials).

### A GLM-HMM reveals dynamic, hidden behavioural states during approach-avoidance conflict

Intermediate responding under conflict (**Fig. 1C**) could reflect a single compromised response rule or alternation between distinct strategies. To distinguish these, we fit a GLM-HMM to trial-by-trial choices (Escola et al., 2011; Ashwood et al., 2022). Within each latent state, choice is modelled as an independent GLM in which state-specific regression weights determine how current predictive cues and recent outcomes are integrated into the probability of a lever press (**Fig. 3A–B**), while the transition matrix defines the persistence of, and switching between, these regimes (**Fig. 3C**). If intermediate conflict responding reflects a single response rule, a one-state model should predict choices as well as a multi-state model. If it reflects alternation between strategies, a multi-state model should predict choices better, particularly on conflict trials (Bolkan et al., 2022).

**Figure 3.**
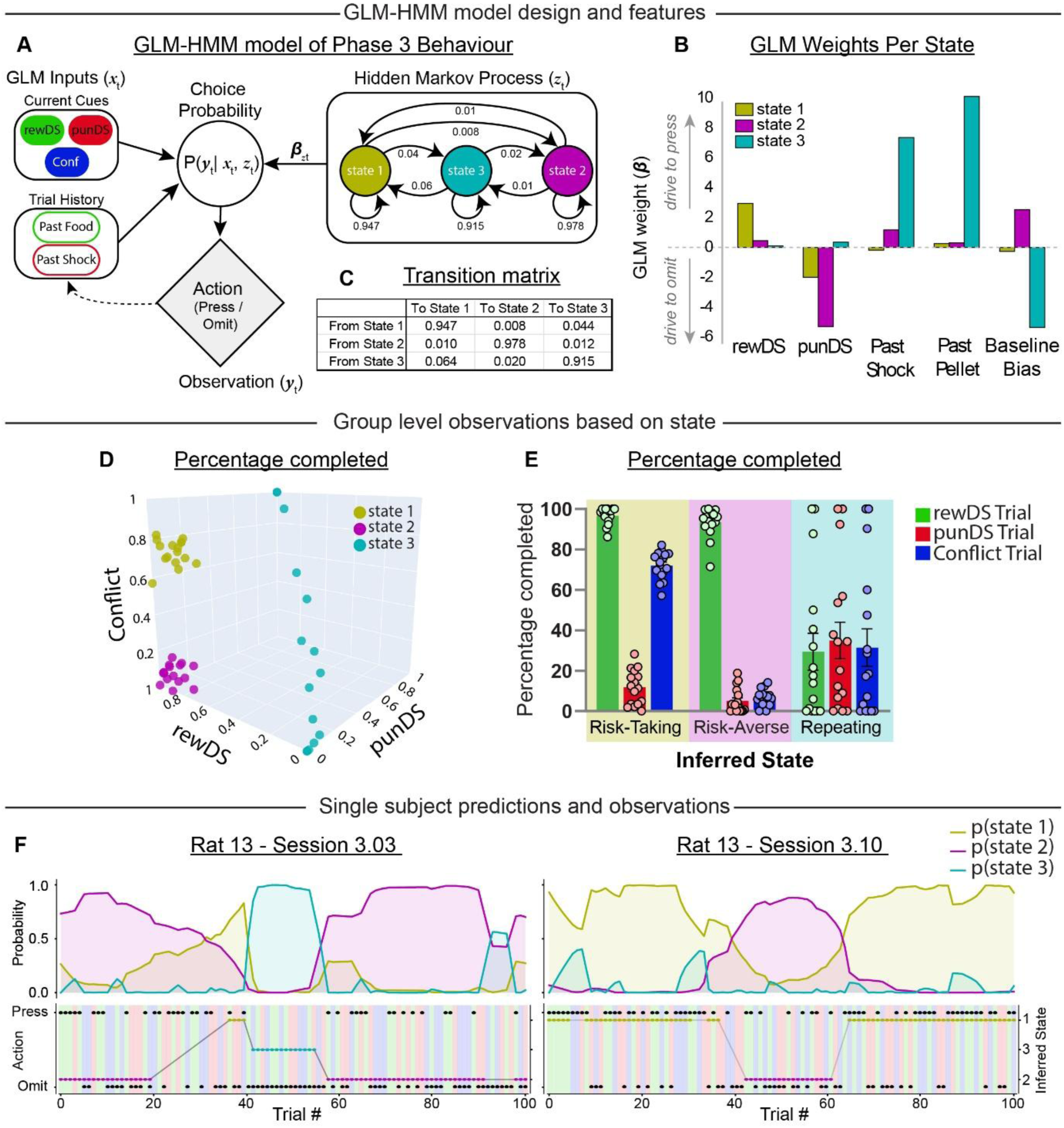
GLM-HMM architecture and latent state dynamics during conflict. **A.** Schematic of the GLM-HMM. On each trial, the probability of a press depends on the cue inputs (rewDS and punDS; both are on during conflict trials) and on the outcome of the previous trial (past pellet, past shock), weighted by the weights of the current latent state. The state persists or switches between trials with the transition probabilities shown, which do not depend on the inputs. **B.** Regression weights mapping the relative influence of external environmental stimuli (rewDS, punDS) and immediate reward or punishment history (Past Pellet, Past Shock) alongside a Baseline Bias parameter for each of the three inferred hidden states. **C.** Table displaying the transition probabilities of the group model remaining in or moving between the three states from trial t to trial t+1. **D.** Response profile of each inferred state. Each point is one rat’s mean response rate on rewDS, punDS and conflict trials assigned to that state. **E.** State-isolated response profiles. Mean (± SEM; n=16) response rates on rewDS, punDS and conflict trials in State 1 (Risk-taking), State 2 (Risk-averse) and State 3 (Repeating). Response rates depended on the interaction between state and trial type (two-way RM ANOVA, State × Cue: F(2.7,41.2) = 725.9, p < 0.0001). In State 1, conflict responding was lower than rewDS (p < 0.0001) and higher than punDS responding (p < 0.0001). In State 2, conflict responding did not differ from punDS (p = 0.63) and was lower than rewDS responding (p < 0.0001). In State 3, punDS responding slightly exceeded rewDS responding (p = 0.0051), and conflict responding did not differ from either (p ≥ 0.12). Šídák-corrected comparisons. **F.** Representative sessions from one rat. Top: posterior probability of each state across trials. Bottom: trial type (background colour), choice (press or omission) and inferred state.

#### Selection of a three-state model

We first fit one-to five-state models to the full behavioural dataset from Phase 3. We validated all models by leave-one-rat-out cross-validation, refitting on 15 rats and evaluating on the held-out rat. Held-out accuracy rose from 77.0 ± 0.9% (single-state) to 83.1 ± 1.0% for the three-state model (**Fig. S2B**). Critically, the advantage of the latent-state model was cue-specific and greatest on conflict trials, where the three-state GLM-HMM predicted held-out choices substantially better than a single-state GLM (74.7 ± 1.9% vs 65.6 ± 1.4%; 16/16 rats; p = 0.0003). A comparable advantage was observed on rewDS trials (+8.5 pp; 14/16 rats; p = 0.002), whereas the two models were indistinguishable on punDS trials (+0.6 pp; 4/16 rats; p = 0.49), consistent with uniformly suppressed responding leaving little latent variance to explain (**Fig. S2A**).

Accuracy, AIC and BIC improved further with a fourth state (**Fig. S2B–D**), so we asked whether the four-state solution contained a new strategy. The Risk-taking and Risk-averse states were recovered almost unchanged (cosine similarity of weights 0.997 and 0.999; **Fig. S2E**). The additional state came from the Repeating state: 70.6% of Repeating trials were assigned to one four-state state (Repeating 1) and 27.0% to the other (Repeating 2; **Fig. S2F**). Rats rarely pressed in Repeating 1, and pressed on most trials of every type in Repeating 2 (**Fig. S2H**). By weight similarity, Repeating 2 most resembled the Risk-averse state (cosine 0.929), because both combine a large positive bias with a negative punDS weight. In Repeating 2, however, the bias (+4.4) offset the punDS weight (−4.4), so most punDS trials were still responded. The fourth state therefore split the Repeating state into high- and low-responding variants. It did not add a strategy that resolved conflict differently from the two on-task states, and we retained the three-state model. A similar choice was made by Venditto et al. (2024), where a fourth state improved fit but split idiosyncratically across animals.

#### Two on-task states resolve conflict in opposing ways

The state-specific regression weights revealed two ’on-task’, stimulus-driven states (States 1 and 2) and a single ’off-task’ state (State 3; **Fig. 3B**). Response rates depended on the interaction between state and trial type (n = 16; **Fig. 3D, E**). In both on-task states rats responded on almost all rewDS trials (State 1: 96.7%; State 2: 93.5%; p = 0.17) and on few punDS trials (11.8% and 5.0%). The two states diverged in how they resolved conflict. In the Risk-taking state (State 1), a strong positive rewDS weight (+2.92) outweighed a moderate negative punDS weight (−1.98), with little influence of trial history (past pellet +0.26, past shock −0.18; **Fig. 3B**). In this state rats responded on most conflict trials (72.0%), far more than on punDS trials (p < 0.0001) but less than on rewDS trials (p < 0.0001); conflict was therefore treated predominantly, though not entirely, as a reward trial. In the Risk-averse state (State 2), a strong negative punDS weight (−5.27) acted against a high baseline bias to respond (+2.51; **Fig. 3B**), and conflict responding (6.9%) was suppressed to the level of punDS responding (5.0%; p = 0.63). Conflict responding differed between the two on-task states (72.0% vs 6.9%; p < 0.0001; Fig. 3E). Thus, the same conflict cue was predominantly approached in one state and treated as a punishment cue in the other.

#### An off-task state implements cue-independent choice perseveration

In contrast to these cue-dependent strategies, the Repeating state (State 3) operated entirely off-task. Weights for the current sensory cues were more than an order of magnitude smaller than the corresponding weights in the on-task states (rewDS +0.10 vs +2.92 in State 1; punDS +0.35 vs −5.27 in State 2), while trial history dominated, with large positive weights for both past pellet (+10.04) and past shock (+7.31; **Fig. 3B**). Because a lever press in this task always produced an outcome, these two regressors sum exactly to whether the animal responded on the preceding trial; their large, similarly-signed weights therefore identify this state as implementing choice perseveration rather than outcome-specific learning. The resulting emission function is near-deterministic and essentially cue-independent: p(press) = 0.99 after a pellet, 0.89 after a shock, versus 0.005 following an omission. This accounts for the bimodal response rates across animals in this state (**Fig. 3D, E**): because the state perpetuates whatever action the animal is already performing, individual rats show near-total responding or near-total omission while occupying it. At the group level, responding in the Repeating state was low and no longer cue-appropriate (rewDS 29.4%, punDS 34.9%, conflict 31.4%). punDS responding slightly exceeded rewDS (p = 0.0025) and conflict responding (p = 0.023), and conflict and rewDS responding did not differ (p = 0.08).

#### States alternate within sessions

Continuous posterior decoding showed that rats did move between states within a session (**Fig. 3F**). States were nonetheless persistent, with high self-transition probabilities (**Fig. 3C**). Occupancy of each state varied considerably between rats (**Fig. S3A**). The Risk-averse state was the most persistent (median 25.5 trials), the Risk-taking state was intermediate (median 11 trials), and the Repeating state median was 8 trials (**Fig. S3B**).

Across 233 included sessions, 76.4% contained at least one state switch (median 2 per session; **Fig. S3C**). Every rat had at least one session in which both on-task states were expressed (33.9% of all sessions; **Fig. S3D**). Most switches were between an on-task state and the Repeating state. Changes between the two on-task states were less frequent and occurred more often directly (83) than via the Repeating state (45). Across Phase 3, Risk-taking occupancy increased (+1.7% per session) and Risk-averse occupancy decreased (−1.9% per session). Repeating occupancy did not change (+0.2% per session; **Fig. S3F**). This shift paralleled the rise in conflict responding across Phase 3 (Fig. 1C).

Because shock intensity was titrated between sessions, we asked whether state occupancy tracked intensity. Within rats, intensity was unrelated to session number (median r = 0.06). Controlling for session, within-rat intensity was not significantly related to occupancy of any state (**Fig. S3G**): Risk-taking: −7.9% per 0.05 mA; Risk-averse: +0.6%; Repeating: +7.2%. Session-to-session changes in intensity also did not predict changes in occupancy (all p ≥ 0.09). State occupancy therefore shifted with training but did not track shock intensity, and much of the variation in state occurred within sessions, where shock intensity was held constant.

### Anterior insular cortex flexibly encodes conflict based on the active latent behavioural state

To determine whether aIC activity statically tracks physical task stimuli or flexibly aligns with the rat’s latent behavioural state, trial-by-trial GLM-HMM state assignments (posterior confidence >= 0.65) were mapped onto aIC photometry recordings. Analysis was restricted to the two primary stimulus-driven states: Risk-taking (State 1) and Risk-averse (State 2), because in State 3 there was minimal within-subject variation between responding and omitting for sufficient comparisons.

We next examined aIC activity on conflict trials across the inferred states, restricting each comparison to rats with at least five trials of a given state × trial-type × choice combination. Conflict-trial activity differed between the Risk-taking and Risk-averse states in specific epochs (**Table 2**). On responded conflict trials, cue-period activity rose above baseline in the Risk-taking state earlier during this epoch (∼5s) but in the Risk-averse state only at the end of the cue. However, the two states did not differ significantly at any time point (**Fig. 4A**). On omitted conflict trials, activity rose above baseline shortly after cue onset only in the Risk-averse state. The states differed for most of the cue period (**Fig. 4B**). Around the conflict lever press, activity rose above baseline in both states and did not differ between them (**Fig. 4C**).

**Figure 4.**
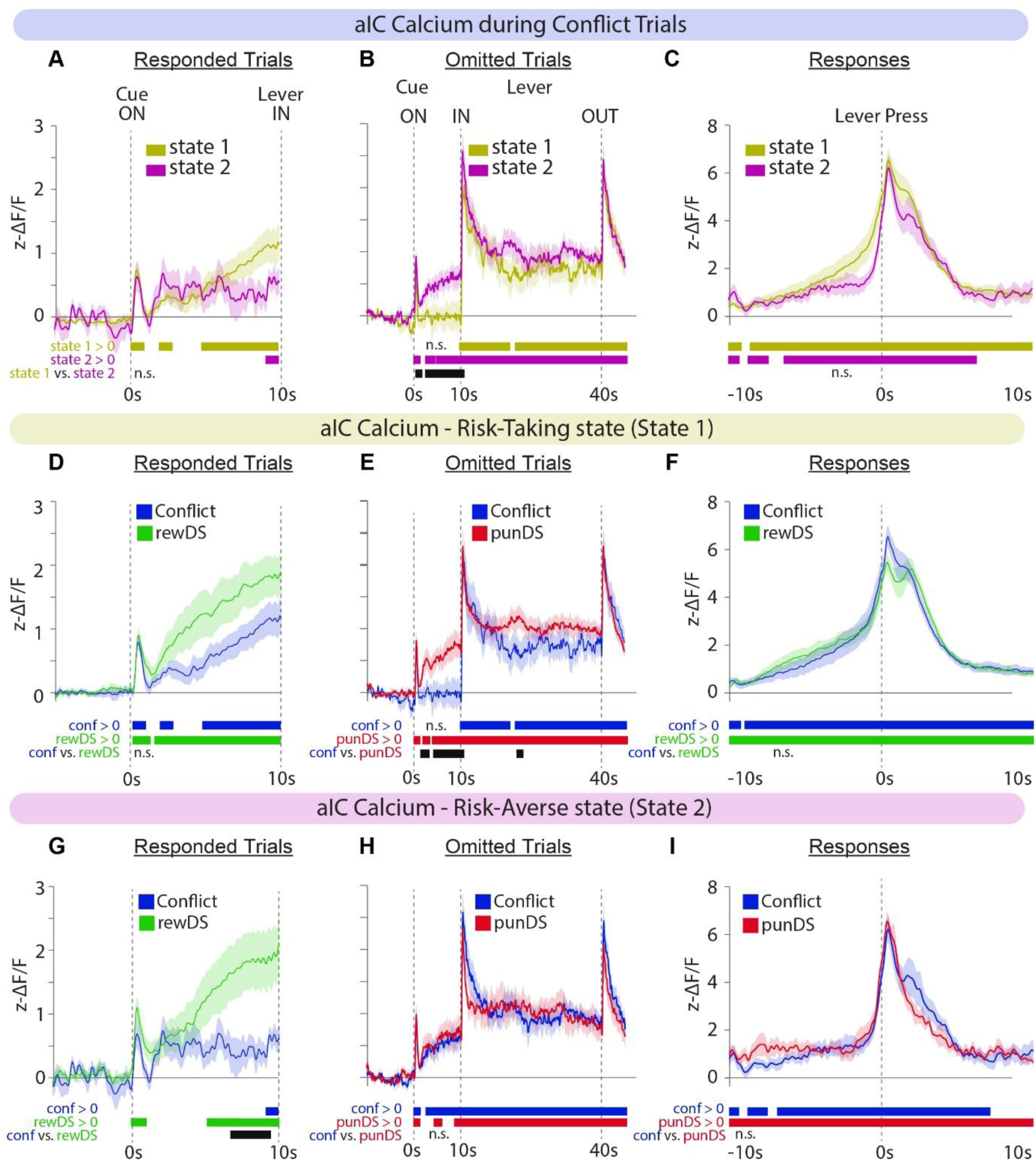
State-dependent modulation of aIC activity during conflict. A-C. Traces depicting aIC activity during conflict trials in Phase 3 show that activity during the different trial epochs is state-dependent. Traces are depicted for the cue period during responded trials (**A**; s1 n=13, s2 n=8), omitted trials (**B**; s1 n=13, s2 n=13), or the activity surrounding choice response (**C**; s1 n=13, s2 n=8). **D-F.** Traces depicting aIC activity during Phase 3 in state 1 showing that, when the rat is in state 1, activity during conflict resembles rewDS and not punDS. Traces are comparing responded rewDS and conflict trials (**D**; rewDS n=14, conflict n=13), omitted punDS and conflict trials (**E**; s1 punDS n=13, conflict n=13), and responses during rewDS and conflict trials (**F**; rewDS n=14, conflict n=13). **G-I.** Traces depicting aIC activity during Phase 3 in state 2 showing that, when the rat is in state 2, activity during conflict resembles punDS and not rewDS. Traces are comparing responded rewDS and conflict trials (**G**; rewDS n=14, conflict n=8), omitted punDS and conflict trials (**H**; punDS n=12, conflict n=13), and responses during punDS and conflict trials (**I**; punDS n=6, conflict n=8). For all trace graphs, the bars underneath represent the time along the trace where the statistical tests (bootstrapped CI, permutation tests) were significant. *Significant time windows are reported in Table 2. n = rats with ≥ 5 trials per condition*.

We then compared conflict trials with rewDS and punDS trials within the on-task states. In the Risk-taking state, responded conflict and rewDS trials did not differ at any time point, either during the cue (**Fig. 4D**) or around the lever press (**Fig. 4F**). On omitted trials, punDS activity rose shortly after cue onset, whereas conflict activity did not exceed baseline until lever insertion, so that the two differed during the cue period (**Fig. 4E**). In the Risk-averse state, cue-period activity on responded conflict trials was lower than on rewDS trials late in the cue (**Fig. 4G**), reversing the pattern seen in the Risk-taking state, where the two did not differ. Omitted conflict and punDS trials did not differ at any time point (**Fig. 4H**). Around the lever press, conflict and punDS activity did not differ (**Fig. 4I**). Thus, in all three epochs, conflict-trial activity resembled rewDS activity in the Risk-taking state and punDS activity in the Risk-averse state. Direct comparisons between states reached significance only on omitted trials (**Fig. 4B**). The state dependence of conflict encoding therefore rests chiefly on the within-state comparisons, and on the choice-prediction analysis below, which tests the between-state difference directly.

#### The relationship between aIC activity and choice follows the latent state

Fig. 4 showed that, on conflict trials, aIC activity resembled rewDS trials in the Risk-taking state and punDS trials in the Risk-averse state. We next asked whether the relationship between aIC activity and the rat’s subsequent choice followed the same pattern. To isolate information not already captured by behaviour, cue-period aIC activity was related to choice beyond the trial-by-trial prediction of the fixed GLM-HMM (**Fig. S4**). In both on-task states, higher aIC activity accompanied the cue-appropriate action: responding on rewDS trials (Risk-taking: odds ratio [OR] = 1.42 per SD; Risk-averse: OR = 1.83; both p < 0.001) and withholding on punDS trials for the Risk-taking state (OR = 0.75, p < 0.001), but not for the Risk-averse state (OR = 0.87, p = 0.16; **Fig. S4B**). On conflict trials, the relationship depended on state. In the Risk-taking state, higher aIC predicted responding (OR = 1.38, p = 0.005), indistinguishable from rewDS trials (p = 0.79) and distinct from punDS trials (p < 0.001; **Fig. S4C**). In the Risk-averse state, the conflict relationship was negative (OR = 0.86), indistinguishable from punDS trials (p = 0.99) and distinct from rewDS trials (p < 0.001; **Fig. S4D**), although it did not differ significantly from zero on its own (p = 0.28). Conflict relationships differed between the two states (p = 0.022; permutation p ≤ 0.001), and no relationship was detected in the Repeating state (all p > 0.18). These relationships reflected genuine trial-level information: cue-period aIC improved held-out prediction beyond a within-cue shuffle (permutation p ≤ 0.001; 12 of 14 rats with cue-specific slopes, p = 0.008), whereas pre-trial activity did not (p = 0.78; **Fig. S4A**). Thus, on conflict trials, aIC activity related to choice as on rewDS trials in the Risk-taking state, and as on punDS trials in the Risk-averse state.

Together, these data show that aIC activity on conflict trials did not take an intermediate value between activity on reward and punishment trials. It resembled activity on rewDS trials in the Risk-taking state and on punDS trials in the Risk-averse state, and its relationship with choice followed the same pattern.

### Acute systemic alcohol alters latent behavioural state occupancy

To determine how pharmacological modulation affects latent state expression and aIC encoding, rats were given acute systemic injections of alcohol (saline, 0.5 g/kg and 1.0 g/kg) before Phase 3 sessions. The first 15 minutes of all test sessions were excluded because during this period, alcohol produced a dose-dependent reduction in rewDS responding (**Fig. S5A**), indicating a general performance impairment that was absent from the remainder of the session. Consistent with this, conflict responding in the first 15 minutes was reduced rather than increased at the high dose (Fig. S5A), and state occupancy was dominated by the Repeating state at the expense of both cue-driven states (**Fig. S5B**). Because responding under conflict can only be meaningfully interpreted when the capacity to respond for reward is intact, all subsequent analyses were restricted to the period after the first 15 minutes. In this period, alcohol altered responding in a trial-type-specific manner (two-way RM ANOVA, Cue × Dose: F(2.97, 41.63) = 9.45, p < 0.0001; n = 15; **Fig. 5A**). rewDS responding did not differ significantly between doses (saline 87.3%, 0.5 g/kg 78.6%, 1.0 g/kg 76.8%; all p ≥ 0.155). At 1.0 g/kg, alcohol increased responding relative to saline on punDS trials (15.6% to 35.2%; p = 0.022) and on conflict trials (46.1% to 62.1%; p = 0.0054), consistent with disinhibition. The 0.5 g/kg dose had no significant effect (punDS p = 0.19; conflict p = 0.44).

**Figure 5.**
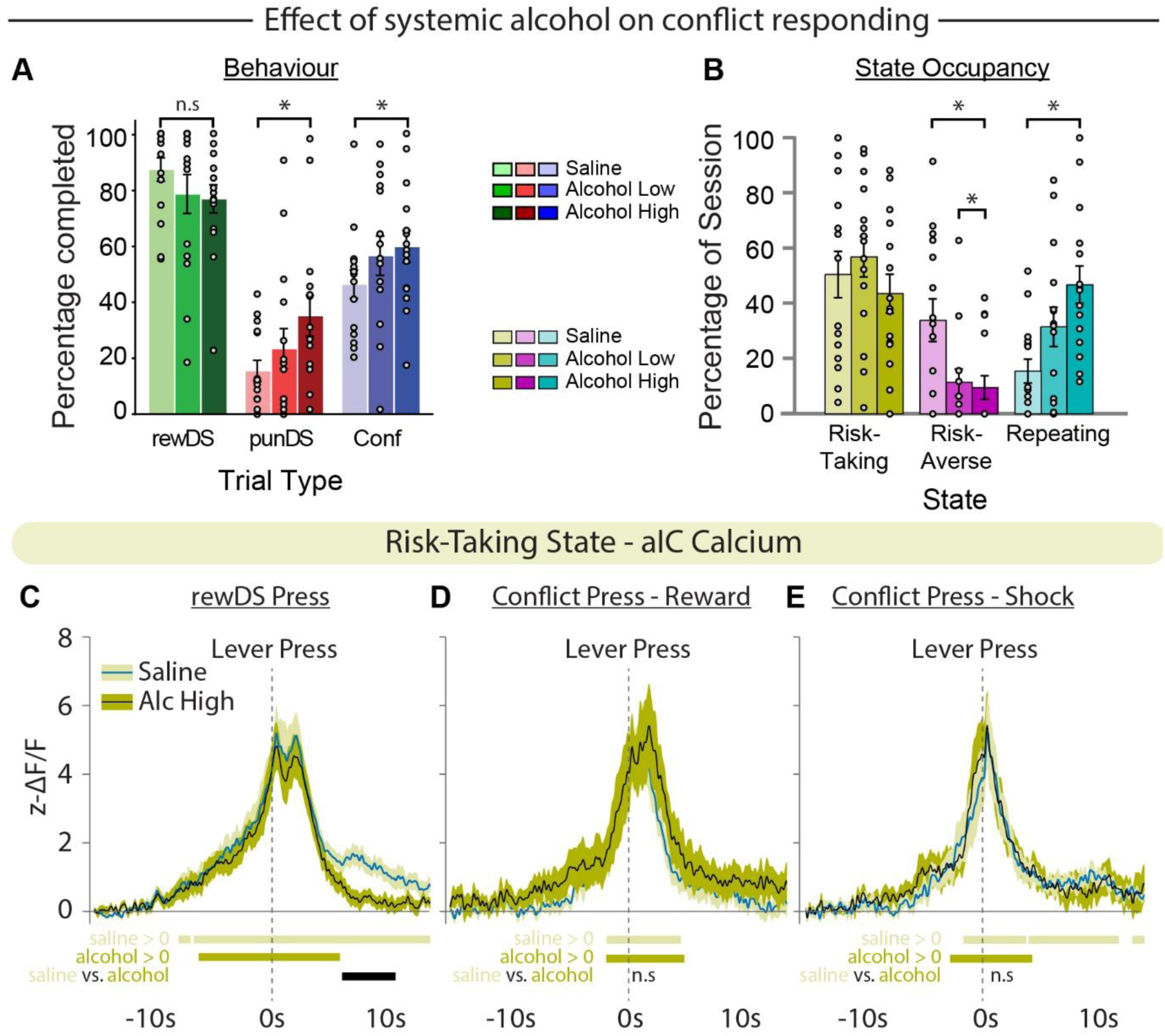
Effect of systemic alcohol on behavioural and state-dependent aIC activity during conflict. **A.** Percentage of trials completed after the first 15 min of the test session (mean ± SEM; points are individual rats, n = 15). Two-way RM ANOVA with Geisser–Greenhouse correction: Cue × Dose, F(2.97, 41.63) = 9.45, p < 0.0001. Saline vs 1.0 g/kg: punDS p = 0.022, conflict p = 0.0054; all other comparisons p ≥ 0.155 (Holm–Šídák). **B.** State occupancy: % of classified trials; mean ± SEM; points are individual rats, n = 15. Two-way RM ANOVA, State × Dose: F(3.36, 47.02) = 4.87, p = 0.0037. Risk-averse: saline vs 0.5 g/kg p = 0.043, vs 1.0 g/kg p = 0.015; Repeating: saline vs 1.0 g/kg p = 0.003; all other comparisons p ≥ 0.14 (Holm–Šídák). **C–E.** aIC activity in the Risk-taking state aligned to the lever press after saline or 1.0 g/kg alcohol, for rewDS presses (**C**) and for conflict presses that ended in reward (**D**) or shock (**E**). Significant time windows are reported in Table 3. *, p < 0.05.

We next used the existing three-state model to decode state occupancy in the alcohol test sessions. Rather than uniformly scaling response probabilities, alcohol altered the distribution of latent behavioural states (two-way RM ANOVA, State × Dose: F(3.36, 47.02) = 4.87, p = 0.0037; n = 15). Compared to saline, acute alcohol caused a reduction in occupancy of the Risk-averse state (33.9% to 11.4% at 0.5 g/kg, p = 0.043; to 9.6% at 1.0 g/kg, p = 0.015) and a corresponding increase in the Repeating state (15.5% to 31.6% at 0.5 g/kg, p = 0.14; to 46.8% at 1.0 g/kg, p = 0.003). At 0.5 g/kg, Risk-averse occupancy fell (p = 0.043) although responding did not change significantly on any trial type, so the state analysis detected an effect of the low dose that aggregate response rates did not. Thus, alcohol-induced disinhibition is not driven by an increase in risk-taking (Risk-taking state: saline 50.5%, 0.5 g/kg 57.0%, 1.0 g/kg 43.6%; all p ≥ 0.37), but by an inability to sustain the cue-driven risk-averse state, resulting in a lapse into rule-insensitive, repeating behaviour.

To assess whether alcohol impacted aIC encoding within the functional states, aIC activity during the Risk-taking state was compared across saline and alcohol conditions. Because alcohol severely diminished Risk-averse occupancy, analysis was restricted to the predominant Risk-taking state. In the Risk-taking state, alcohol (1.0 g/kg) did not alter pre-press ramping or the press-evoked response on rewDS trials, but it reduced the late post-reward component of the response (**Fig. 5C**; **Table 3**). Activity around conflict presses that ended in reward (**Fig. 5D**) or shock (**Fig. 5E**) did not differ between saline and alcohol at any time point.

### Anterior insular cortex encoding of conflict requires instrumental agency

To determine whether aIC engagement in this task requires instrumental agency, we recorded aIC in rats tested in a yoked Pavlovian design of the task. These rats received the exact sequence of trial type, duration, and outcome generated by a paired Operant counterpart, but everything was delivered independent of their behaviour, lever presses had no consequence (**Fig. 6A**). For analysis of the photometry data, only the specific rat and sessions from the Operant rats which were used for the yoking were included in the analysis.

**Figure 6.**
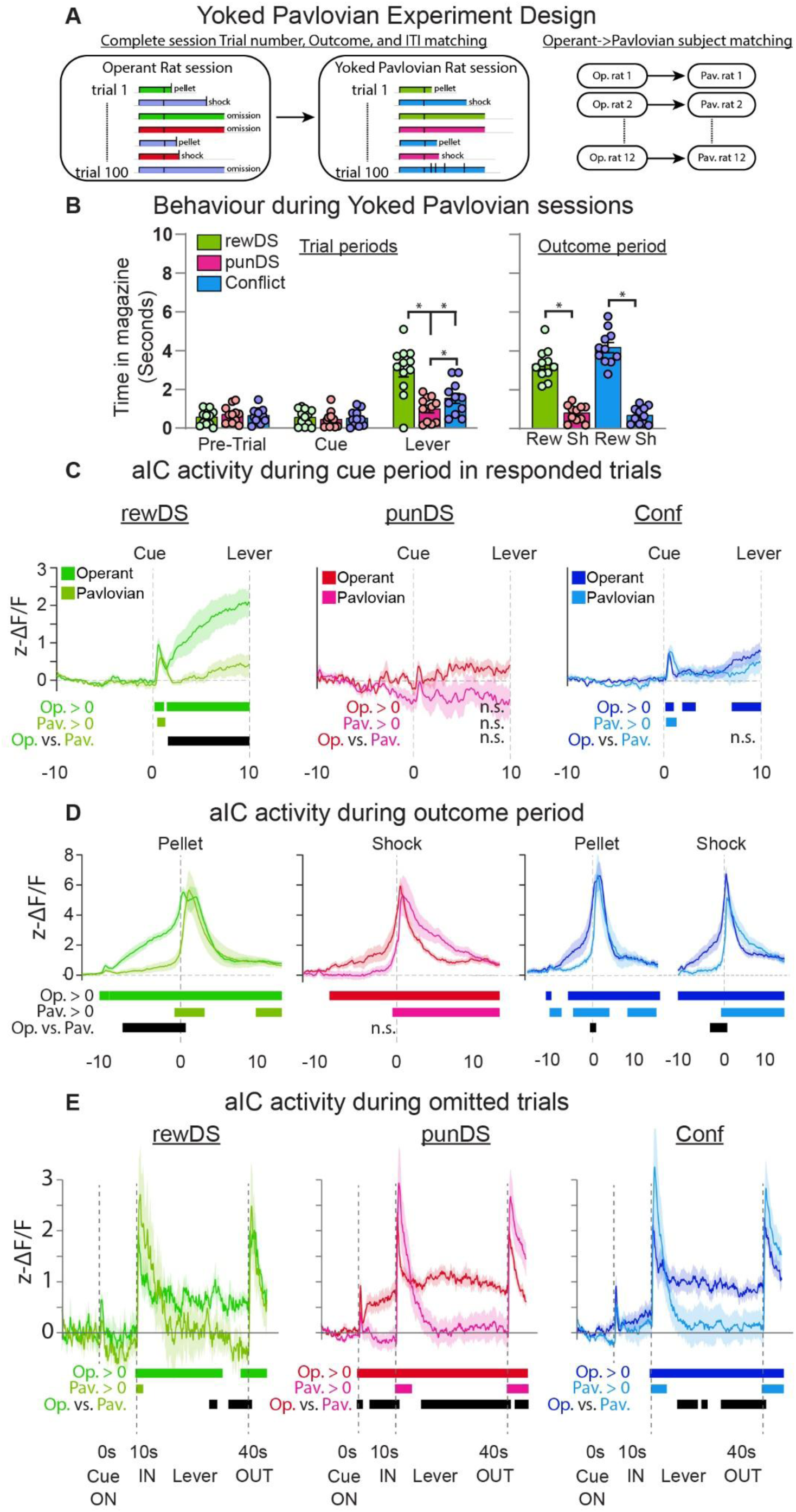
aIC activity in Operant and yoked Pavlovian rats during conflict training. **A.** Description of the Yoking procedure, where the specific trial types, timing of trial start and end, outcomes, and ITIs are mapped from a matched Operant cohort rat to a Pavlovian Cohort rat, throughout the entire experiment. **B.** Behavioural observations (time in magazine) during the yoked Pavlovian task in Phase 3. Lever-period time in magazine was significantly higher for Conflict trials compared to punDS trials, but significantly lower than rewDS trials (mixed-effects analysis: Cue × Epoch interaction, F(1.3, 12.2) = 43.71, p < 0.0001; post-hoc Conflict vs. punDS: p = 0.003, rewDS vs. Conflict: p = 0.0001). During the 10 s after outcome delivery, magazine time was higher after rewarded than after shocked conflict trials and higher after rewarded conflict trials than after rewDS rewards, and did not differ between shocked conflict and punDS trials (mixed-effects model: main effect of trial type, F(2.03, 20.34) = 130.9, p < 0.0001; Šídák: conflict-reward vs conflict-shock p < 0.0001, conflict-reward vs rewDS p = 0.012, conflict-shock vs punDS p = 0.86). **C.** Traces depicting aIC activity during the cue period (cue on = 0s) in completed trials in Phase 3. rewDS trials: Pav n=9; Op n=9. punDS trials: Pav n=7; Op n=7. Conf trials: Pav n=9; Op n=9. **D.** Traces depicting aIC activity surrounding lever press (Operant) or outcome delivery (Pavlovian) in Phase 3. rewDS trials: Pav n=9; Op n=9. punDS trials: Pav n=7; Op n=7. Conf trials: Pav n=9; Op n=9. **E.** Traces depicting aIC activity during the entire trial period (cue on = 0s) in non-completed trials in Phase 3. rewDS trials: Pav n=5; Op n=6. punDS trials: Pav n=9; Op n=9. Conf trials: Pav n=9; Op n=9. *Significant time windows are reported in Table 4.* *, p<0.05. Pav, Pavlovian; Op, Operant. N reflects number of rats that met the threshold of at least 5 trials per condition.

Pavlovian rats successfully acquired cue contingencies in Phase 2 (**Fig. S6A**). During the cue, magazine time was slightly lower on punDS than rewDS trials (p = 0.027), and the difference was larger during the lever period (p < 0.0001). Time in magazine was also significantly higher following reward delivery compared to foot-shock. This discrimination persisted into Phase 3, where lever-period time in magazine after rewarded Conflict trials was higher than after rewDS rewards (p = 0.012), while after shocked Conflict trials it did not differ from punDS trials (p = 0.86) (**Fig. 6B**). Finally, focusing on the 10s period after outcome delivery revealed that time in magazine during rewarded conflict trials was statistically higher than rewDS trials, while shocked conflict trials were indistinguishable from punDS trials.

Despite having learned the task contingencies, Pavlovian rats showed markedly different task-evoked aIC activity from Operant rats. On responded rewDS trials, cue-period activity rose above baseline in Operant rats, but only transiently in Pavlovian rats, and the groups differed significantly (**Fig. 6C**). On responded punDS trials, activity did not exceed baseline in either group, and the groups did not differ. On completed Conflict trials (i.e. Operant rat responded), activity exceeded baseline for part of the cue period, but the groups did not differ (**Table 4**). Pavlovian rats showed robust responses to the delivery of both reward and shock (**Fig. 6D**), confirming that aIC signals were readily detectable in this cohort. The groups differed mainly before the outcome. On rewDS trials, Operant activity was elevated prior to the lever press and differed significantly from the Pavlovian cohort; on punDS trials the groups did not differ, and there were some differences between the groups in the conflict trials (**Fig. 6D**).

Crucially, isolation of non-completed punDS and Conflict trials (i.e. Operant rat did not respond) showed that avoidance-related aIC activity depended on instrumental control. In the Pavlovian cohort, the presence of the lever is a necessary but unreliable predictor of footshock in punDS and conflict trials. On non-completed punDS trials, activity in Pavlovian rats exceeded baseline only transiently after lever insertion and lever retraction. Activity was less in Pavlovian rats compared to Operant rats during the cue period and for most of the lever period (**Fig. 6E**). In conflict trials we observed a similar pattern: activity in Pavlovian rats was increased above baseline only transiently, and the groups differed intermittently. On non-completed rewDS trials (Operant n=6; Pavlovian n=5), the groups differed only late in the lever period.

A similar dissociation was already present in Phase 2 (**Fig. S6; Supplementary Table S1**). On completed trials, cue-period activity exceeded baseline in Operant but not Pavlovian rats (**Fig. S6B**). Activity to the outcomes themselves was intact in Pavlovian rats (**Fig. S6C**). On non-completed punDS trials, Pavlovian rats did show activity above baseline during much of the lever period, but it was lower than in Operant rats (**Fig. S6D**). Non-completed rewDS trials did not differ between groups.

Collectively, these data demonstrate that the characteristic neural signatures of approach-avoidance conflict in aIC are not merely sensory responses to ambiguous cues or unpredicted outcomes. These results indicate that the sustained aIC activity on omitted trials in Operant rats depended on the opportunity to withhold a goal-directed action, not on the cue and lever presentation alone.

## Discussion

We asked whether decision making under approach–avoidance conflict reflects a single policy reflecting summation of potential positive and negative outcomes, or alternation between discrete strategies. We found that choice in conflict was better described by a latent state model, whereby each state serves as a distinct GLM, and action selection in conflict trials was resolved in opposing ways depending on the state. We next found that aIC activity on conflict trials was modulated depending on state, and the relationship between cue-period aIC activity and choice differed between the two ‘on-task’ states. We next showed that acute alcohol increased both punished and conflict responding and applying the latent-state strategy revealed that this disinhibition was explained by reduced occupancy of the Risk-averse state rather than increased Risk-taking. Finally, we show that the aIC signals that accompanied pressing or withholding were largely absent in yoked rats that received the same cues and outcomes without control over them, demonstrating that volitional action is necessary for recruitment of aIC during specific aspects related to decision-making in conflict.

### Intermediate conflict responding is a mixture of strategies

Integration accounts explain conflict resolution as the summation of the values predicted by the reward and punishment cues, applied on every trial (Miller, 1944; Rangel et al., 2008; Levy and Glimcher, 2012). Common-scale representations of this kind have been described from prefrontal cortex to subcortical regions (Basten et al., 2010; Bartra et al., 2013; Hulsman et al., 2024; Gordon et al., 2026). Summation predicts graded responding that settles at a point of balance between the competing demands (McNally, 2021). However, only one response can be expressed at a time, and behaviour under motivational competition is typically bistable, with one tendency dominating the other (Choi and Kim, 2010; Schlund et al., 2016; Verharen et al., 2019; McNally, 2021; Choi et al., 2022). A binary choice on each trial does not by itself distinguish integration from selection, because a single combined value would also produce binary choices at a constant, intermediate probability. In the model, this probability took one of two values depending on the state. It was high in the Risk-taking state and low in the Risk-averse state, and each state persisted over tens of trials, as reported for perceptual decisions and reversal learning in mice, and for multi-step reward learning in rats (Ashwood et al., 2022; Bolkan et al., 2022; Le et al., 2023; Venditto et al., 2024). The intermediate rate seen before classification (**Fig. 1C**) therefore reflects averaging across states, and neither on-task state responds at that rate. This does not mean that rats failed to attribute valence to the cues. Within each state, choice was an additive combination of inputs, and the states differed in the weight attributed to each cue. Latent states inferred from behaviour alone are hypotheses that require independent validation (Versteijnen and Heimel, 2026). It is therefore important that the states also differed in aIC activity and in the relationship between aIC activity and choice (**Fig. 4, S4**), neither of which was used to fit the model. Behavioural states inferred in the same way have predicted response times and OFC encoding in rats (Venditto et al., 2024).

Related to the Repeating states, states dominated by choice history or bias are common in GLM-HMM analyses and are usually interpreted as task disengagement (Ashwood et al., 2022; Bolkan et al., 2022; Oostland et al., 2026; Venditto et al., 2024). The four-state model split this state into high- and low-responding variants (**Fig. S2**). Bolkan et al. (2022) reported a similar split of their choice-history state, which added little interpretive insight. Additional states often describe finer structure within a state rather than distinct strategies (Pohle et al., 2017; Versteijnen and Heimel, 2026).

### State dynamics and individual differences

States were persistent but not fixed. Dwell times were comparable to those of engaged and biased states in mice (Ashwood et al., 2022; Bolkan et al., 2022). Persistence of this kind is predicted by selection through positive feedback or mutual inhibition, but not by summation (McNally, 2021). All rats switched states within sessions, so the states do not reflect different strategies on different days (Ashwood et al., 2022). Within-session variation accounted for about half of the variance in membership of the Risk-taking and Repeating states, and cannot reflect shock intensity, which changed only between sessions. Changes in motivational state, such as satiety, are one candidate (Bolkan et al., 2022; Gordon et al., 2026; Venditto et al., 2024). Because transition probabilities in our model were stationary, the analysis describes when switches occurred but not what caused them. Models in which transitions depend on trial inputs (Calhoun et al., 2019; Mohammadi et al., 2025) could test whether recent outcomes, such as a shock on a conflict trial, make a switch more likely. In mice, such a model showed that past rewards predicted switches into disengaged states (Mohammadi et al., 2025). In humans, a shock on the previous trial slowed subsequent approach to a conflict cue (Chen et al., 2026).

Our data suggests that how readily, and for how long, a rat adopts a Risk-averse strategy describes a potential computational mechanism describing punishment resistance. Occupancy of the Risk-averse state varied more between rats than the others. Punishment sensitivity is often bimodal in rats (Pelloux et al., 2007; Marchant et al., 2018; Jean-Richard-Dit-Bressel et al., 2019) and humans (Jean-Richard-Dit-Bressel et al., 2021), and rats show stable individual differences in risk preference (Simon et al., 2011). Crucially, when occupying the Risk-taking state, rats still withheld on most punDS trials, so infrequent use of the Risk-averse state cannot be explained by failure to learn punishment contingencies (Jean-Richard-Dit-Bressel et al., 2019; Jean-Richard-Dit-Bressel et al., 2021). Across Phase 3, Risk-taking occupancy increased and Risk-averse occupancy decreased, in parallel with the rise in conflict responding. In classic runway studies, shock intensity determined which tendency dominated, and behaviour was least stable at intermediate intensities (Miller, 1944; McNally, 2021). Within rats, shock intensity was not significantly related to occupancy of any state. Because intensity varied over a narrow range and was titrated in response to behaviour, we interpret this null result cautiously. Testing whether punishment magnitude shifts state occupancy, within-state weights, or both, would either require intensities set independently of behaviour, or entered as a covariate on the model parameters (Kirchherr et al., 2023; Versteijnen and Heimel, 2026).

### aIC activity on conflict trials follows the active state

Within each state, conflict activity resembled activity on the certain trial type (rewDS, punDS) that dominated that state’s policy. Across the task, aIC activity accompanied the action appropriate to the current cue, whichever direction that action took. This pattern was also evident in conflict trials. We show that aIC activity followed the action dictated by the active state. If these signals reflect the selection of an action strategy, they should depend on the capacity to act, and the yoked Pavlovian experiment tested this. We show that aIC does respond to reward and shock, however there were key differences in this activity compared to the operant group. These components therefore depended on control over the outcome rather than on the cues and outcomes themselves. This is consistent with a role for the insula in retrieving outcome values that guide goal-directed action (Parkes and Balleine, 2013). It is also consistent with insular involvement in the sense of agency (Farrer and Frith, 2002) and in the perceived controllability of aversive events (Salomons et al., 2004). The activity on omitted trials is the most informative result, because no action was performed. In operant rats, sustained activity lasted for the entire trial in which the rat withheld a response. In yoked rats however, the same cue and lever presentation evoked transients around trial start, lever insertion, and lever retraction, without the sustained activation seen in the operant rats. Together, these data indicate that aIC activity is related to decision-making and action selection, even when the action is withholding a response.

#### Alcohol shifts occupancy away from the Risk-averse state

We found that acute alcohol increased responding on both punDS and conflict trials without significantly changing rewDS responding. This reproduces the well-established anti-punishment effect of alcohol and anxiolytics in conflict procedures (Geller and Seifter, 1960; Vogel et al., 1971). Our latent-state analysis suggests how this anti-punishment effect arises. We show that alcohol reduced occupancy of the Risk-averse state and increased occupancy of the Repeating state, while Risk-taking occupancy did not increase. Therefore, increased punished responding under alcohol does not reflect adoption of a Risk-taking strategy, rather it reflects a failure to sustain the Risk-averse strategy and a lapse into history-driven responding. This shows that state-based analyses can detect drug effects on strategy that aggregate response rates miss. One implication of this interpretation is that acute alcohol may promote drinking despite negative consequences by weakening the strategy that implements punishment sensitivity, rather than by increasing the value of reward.

#### Limitations

Our fibers were placed in aIC, however at these anteroposterior levels aIC borders lateral orbitofrontal cortex (lOFC), so we cannot exclude a contribution from adjacent tissue. This matters for interpretation, because properties such as dependence on latent state are also characteristic of lOFC (Wilson et al., 2014; Schuck et al., 2016) and OFC encoding of outcome value and reward is modulated by HMM-inferred behavioural state (Venditto et al., 2024). aIC and lOFC are densely and reciprocally connected (Gehrlach et al., 2020), and the state-dependent signals we recorded may reflect computations shared across this region rather than computations unique to aIC.

The state dependence of conflict-trial activity in Fig. 4 rests mainly on comparisons within each state. Only eight rats had enough responded conflict trials in the Risk-averse state. Comparisons between states involved partly different rats, and the absence of a significant difference is not evidence of equivalence. The direct test of a between-state difference is therefore the choice-prediction analysis (**Fig. S4**), which used all trials from 14 rats. Finally, only male rats were tested. Female rats have been reported to be more risk-averse than males in a punished risky-choice task (Orsini et al., 2016). Whether sex affects the occupancy of conflict strategies, their weights, or the aIC signals that accompany them remains to be determined.

## Conclusion

Rats resolved approach–avoidance conflict by alternating between discrete strategies that persisted for tens of trials, rather than by applying a single compromise. Within each strategy, the reward and punishment cues were attributed with different weights, so the same conflict cue was approached under one strategy and avoided under another. aIC activity on conflict trials followed the strategy in force, and its anticipatory and avoidance-related components depended on the capacity to act. Alcohol increased punished responding through loss of the risk-averse strategy rather than through increased risk-taking.

## Acknowledgments

The authors gratefully acknowledge: Dr. Philip Jean-Richard-Dit-Bressel for assistance with photometry analysis and sharing early versions of the scripts we used; Professor Alexander Heimel for advice on the modelling approach; the VUmc Histology Imaging Unit for their support C assistance in whole-slide imaging.

---

**Supplementary Figure 1.** A. Schematic of the surgical approach for fiber photometry recordings. Rats were unilaterally injected with an adeno-associated virus expressing the calcium indicator jGCaMP7f (AAV9-mCaMKIIa-jGCaMP7f) into the anterior insular cortex (aIC), followed by the implantation of an optic fiber above the injection site. **B.** Representative coronal fluorescent micrograph demonstrating jGCaMP7f expression and the optic fiber track terminating just above the aIC. **C,D.** Histological reconstruction of estimated optic fiber tip placements mapped onto standardized coronal atlas sections. Placements are shown for all subjects included in the final analyses for both the Operant cohort (C, n=14) and the yoked Pavlovian cohort (D, n=9). Rats excluded for low expression or fiber placement are not shown.

**Supplementary Figure 2.** Model selection and comparison of the three- and four-state GLM-HMM solutions. A-B. Leave-one-rat-out cross-validation (n = 16 rats; 16 folds, 10 independent random initialisations per fold, 23,126 held-out trials). Models were refit on 15 rats and evaluated on the excluded animal, with choices predicted one step ahead so that no prediction was conditioned on the choice being predicted. **A.** Held-out accuracy of the three-state GLM-HMM (dark) and a single-state GLM baseline (grey), separated by trial type (mean ± SEM across rats; dotted line, chance). The latent-state model conferred its largest advantage on conflict trials, where it outperformed the single-state model in all 16 rats (rewDS: 87.9±1.9% vs 79.4±3.7%, t(15) = 3.87, p = 0.002; punDS: 86.3±2.8% vs 85.7±3.0%, t(15) = 0.71, p=0.492; conflict: 74.7±1.9% vs 65.6±1.4%, t(15) = 4.68, p = 0.0003). **B.** Held-out accuracy for models with 1–5 latent states (mean ± SEM across rats). Accuracy rose from 77.0±0.9% (one state) to 83.1±1.0% (three states) and increased only marginally with additional states (4 states: 84.3±1.0%; 5 states: 84.4±1.0%). Akaike (**C**) and Bayesian (**D**) information criteria for models with 1–5 states fitted to the full dataset (16 rats, 23,126 trials). The dashed red line marks the retained three-state model. **E.** Cosine similarity between the GLM weight vectors of each state of the three-state model (rows) and the four-state model (columns); States were matched by optimal assignment; matched pairs are shown in bold. The Risk-taking and Risk-averse states were recovered almost exactly (cosine 0.997 and 0.999). Most Repeating trials were assigned to Repeating 1 (cosine 0.721). The additional state, Repeating 2, had weights most similar to the Risk-averse state (0.929) because of its large positive bias, but rats responded on most trials of every type in this state (**H**). **F.** Correspondence of trial assignments between the two solutions (state assigned when its posterior probability was ≥ 0.65 in each model). Each row shows the percentage of trials assigned to a three-state state (or left unassigned) that were assigned to each four-state state or left unassigned. **G.** GLM weights of the four-state solution (positive values drive pressing, negative values drive omission). **H.** Percentage of rewDS, punDS and conflict trials completed in each state of the four-state solution (mean ± SEM; points are individual rats).

**Supplementary Figure 3.** GLM-HMM Latent state occupancy, persistence and dynamics. **A.** Overall state occupancy in the three-state model (n = 16 rats), expressed as the percentage of confidently classified trials (posterior probability ≥ 0.65) spent in State 1 (Risk-taking), State 2 (Risk-averse) and State 3 (Repeating). Bars show group mean ± SEM; dots represent individual rats, connected by lines. **B.** State persistence, shown as the distribution of dwell times (number of consecutive trials spent in a state). Runs bridged gaps of up to three consecutive unclassified trials when the same state resumed on both sides, and bridged trials were counted towards run length. Runs ending at the session boundary are right-censored. Boxplots show the median and interquartile range, with whiskers extending to 1.5× the interquartile range, overlaid with individual runs pooled across rats. The number of runs (n) and the median run length (med) are shown above each state. **C.** Distribution of state switches per session (233 sessions, 16 rats). A switch was counted between consecutive runs of different states, each at least three trials long. **D.** Switching per rat: for each rat, the number of Phase 3 sessions in which both on-task states were expressed (≥ 5 trials each; black), sessions with other state switches (grey) and sessions without a switch (white). Rats are ordered by the number of sessions containing both on-task states. One rat completed 8 Phase 3 sessions. **E.** Variance in trial-level state membership attributable to differences between rats, between sessions within rats, and within sessions. Shock intensity was constant within each session. **F.** State occupancy across Phase 3 sessions (mean ± SEM; sessions 1–8, n = 16 rats; sessions 9–15, n = 15 rats). Linear mixed model, occupancy ∼ session number: Risk-taking +1.7% per session (95% CI 1.0 to 2.5, p < 0.0001), Risk-averse −1.9% per session (95% CI −2.7 to −1.1, p < 0.0001), Repeating +0.2% per session (95% CI −0.6 to 0.9, p = 0.64). **G.** Effects of within-rat shock intensity (per 0.05 mA) and session number (per 5 sessions) on session occupancy (estimate ± 95% CI). Both were estimated jointly in linear mixed models that also included between-rat intensity and a random intercept for rat. Within-rat intensity: Risk-taking −7.9% (95% CI −16.5 to 0.8, p = 0.07), Risk-averse +0.6% (95% CI −8.4 to 9.6, p = 0.90), Repeating +7.2% (95% CI −1.6 to 16.1, p = 0.11). Session, adjusted for intensity (per 5 sessions): Risk-taking +8.1% (95% CI 4.2 to 12.0, p < 0.0001), Risk-averse −9.5% (95% CI −13.5 to −5.5, p < 0.0001), Repeating +1.3% (95% CI −2.7 to 5.3, p = 0.52). Within rats, intensity was unrelated to session number (median r = 0.06). In a change model, session-to-session changes in intensity did not predict changes in occupancy (Risk-taking p = 0.12, Risk-averse p = 0.09, Repeating p = 0.75).

**Supplementary Figure 4.** Cue-period aIC activity predicts choice in a state-dependent manner. **A.** Improvement in held-out prediction of choice (leave-one-rat-out cross-validated log-likelihood) when aIC activity was added to the fixed GLM-HMM prediction, for pre-trial and cue-period activity. Bars, mean ± SEM across rats; yellow lines, 95th percentile of a null distribution from 1,000 within-cue shuffles of aIC; p values, permutation test. **B** Coefficients relating cue-period aIC activity to the log-odds of responding, beyond the GLM-HMM prediction, for each trial type within each latent state (95% CI, rat-clustered). **C–D** The same coefficients shown as the change in log-odds of responding across the range of aIC activity, comparing conflict trials (dashed) with rewDS and punDS trials in the Risk-taking (**C**) and Risk-averse (**D**) states. Shaded regions, 95% CI. n = 14 rats

**Supplementary Figure 5.** Behavioural and latent state dynamics during the initial phase of acute alcohol intoxication. **A.** Trial-averaged behavioural response rates (rewDS, punDS, and Conflict) during the first 15 minutes of the test sessions (n = 15; two-way RM ANOVA with Geisser–Greenhouse correction, Cue × Dose: F(2.74, 38.31) = 20.52, p < 0.0001). High-dose alcohol (1.0 g/kg) caused a significant reduction in responded rewDS trials (95.7% to 52.1%; p < 0.0001, Holm–Šídák), reflecting transient motor impairment and task disengagement immediately following systemic administration. **B.** GLM-HMM latent state occupancy during the corresponding 15-minute epoch (n = 15; two-way RM ANOVA, State × Dose: F(2.75, 38.56) = 19.03, p < 0.0001). Acute alcohol (1.0 g/kg) significantly reduced occupancy in the stimulus-driven decision-making states (State 1: Risk-taking, 63.8% to 29.5%, p = 0.0031; State 2: Risk-averse, 26.8% to 2.9%, p = 0.022) while inducing a transient increase in the rule-insensitive, disengaged state (State 3: Repeating, 9.4% to 67.6%, p < 0.0001; Holm–Šídák, saline vs 1.0 g/kg). **C–D.** State persistence (dwell times): number of consecutive trials spent in each state in the first 15 minutes (**C**) and after the first 15 minutes (**D**) of the test sessions. Runs were computed within each epoch, so runs in C cannot exceed the number of trials in that epoch (range: 16–21). Boxes show the median and interquartile range; points are individual runs.

**Supplementary Figure 6.** Behavioural anticipation and aIC activity in the yoked Pavlovian cohort during Phase 2. **A.** Time in magazine (s) during the pre-trial, cue and lever periods, and during the 10 s after outcome delivery, on rewDS and punDS trials (Pavlovian n=12). During Phase 2, time was significantly elevated during the 10s lever-availability period for rewDS compared to punDS trials (repeated-measures ANOVA: Cue × Epoch interaction, F(1.2, 13.4) = 31.82, p < 0.0001; post-hoc rewDS vs. punDS: p < 0.0001). Furthermore, magazine time was significantly higher following reward delivery compared to foot-shock (paired t-test: t(11) = 6.91, p < 0.0001). **B-D.** Mean z-scored aIC calcium activity traces for the yoked Pavlovian cohort during Phase 2. Traces are depicted for completed trials during the cue period (**B;** rewDS trials: Pav n=9; Op n=9. punDS trials: Pav n=9; Op n=9.) and outcome delivery (**C;** rewDS trials: Pav n=9; Op n=9. punDS trials: Pav n=9; Op n=9.); or across the entire trial period for non-completed trials (**D;** rewDS trials: Pav n=6; Op n=9. punDS trials: Pav n=9; Op n=9.). Significance bars beneath the traces represent time-windows where statistical tests (bootstrapped confidence intervals or permutation tests) yielded significant deviations from baseline or between conditions. *Significant time windows are reported in Supplementary Table S1.* *, p<0.05. *Pav*, Pavlovian; *Op*, Operant.

