## Supplemental Tables for "Latent state selection, not outcome integration, resolves approach–avoidance conflict"

| Experimental Phase | Figure | Statistical Comparison | Significance time window (Seconds; 0 = Cues turned on / Lever press) |
|---|---|---|---|
| Phase 1 | 2B | BCI: Responded, rewDS (cue) | n.s. |
| | 2F | BCI: Lever press, rewDS | -3.52→6.29; 10.38→10.95; 11.14→15.04 |
| | 2I | BCI: Omitted, rewDS | 10.24→11.31; 19.48→19.98; 40.35→40.91 |
| Phase 2 Early | 2C | BCI: Responded, rewDS (cue) | n.s. |
| | 2C | BCI: Responded, punDS (cue) | n.s. |
| | 2C | PT: Responded (rewDS v punDS) | n.s. |
| | 2G | BCI: Lever press, rewDS | -4.53→4.95 |
| | 2G | BCI: Lever press, punDS | -2.58→7.30; 7.55→8.62; 9.56→10.13 |
| | 2G | PT: Lever press (rewDS v punDS) | n.s. |
| | 2J | BCI: Omitted, rewDS | 10.30→10.99; 40.79→41.42; 42.36→42.99 |
| | 2J | BCI: Omitted, punDS | 10.18→14.51; 16.27→16.78; 18.85→19.35; 20.17→23.94; 24.26→24.70; 24.89→35.82; 36.07→44.75 |
| | 2J | PT: Omitted (rewDS v punDS) | 37.52→37.96; 39.66→40.54 |
| Phase 2 Late | 2C | BCI: Responded, rewDS (cue) | 0.31→0.94; 1.89→10.00 |
| | 2C | BCI: Responded, punDS (cue) | 5.03→5.47; 5.85→10.00 |
| | 2C | PT: Responded (rewDS v punDS) | n.s. |
| | 2G | BCI: Lever press, rewDS | -10.07→-9.44; -8.68→15.04 |
| | 2G | BCI: Lever press, punDS | -9.94→-9.31; -9.19→-8.24; -7.99→13.84 |
| | 2G | PT: Lever press (rewDS v punDS) | 1.82→4.28 |
| | 2J | BCI: Omitted, rewDS | 3.89→4.90 |
| | 2J | BCI: Omitted, punDS | 0.37→0.87; 7.03→7.47; 7.79→8.29; 8.61→9.23; 10.18→45.00 |
| | 2J | PT: Omitted (rewDS v punDS) | 2.32→3.07; 3.33→5.84; 7.10→9.30; 9.67→15.96; 16.40→18.73; 18.91→21.30; 21.93→43.99; 44.18→45.00 |

**Table 1. Time periods (in seconds) where the photometry statistical tests are significant in Figure 2.** *Time 0 refers to cue onset for responded (B, C) and omitted (I, J) trials, and to the lever press for lever-press traces (F, G). On omitted trials the lever was inserted at 10 s and retracted at 40 s. BCI, Bootstrapped Confidence Intervals; PT = Permutation Tests.*

| Experimental Phase | Figure | Statistical Comparison | Significance time window (Seconds; 0 = Cues turned on / Lever press) |
|---|---|---|---|
| Phase 3 Conflict trials | 4A | BCI: Responded conflict (cue), State 1 | 0.31→0.81; 2.07→2.63; 5.09→10.00 |
| | 4A | BCI: Responded conflict (cue), State 2 | 9.23→9.99 |
| | 4A | PT: Responded conflict (cue) (State 1 v State 2) | n.s. |
| | 4B | BCI: Omitted conflict, State 1 | 10.24→20.11; 22.06→22.56; 23.44→24.76; 25.83→27.97; 29.10→33.18; 33.87→45.00 |
| | 4B | BCI: Omitted conflict, State 2 | 0.31→1.00; 3.01→4.14; 5.27→6.34; 6.47→8.48; 9.55→24.13; 24.45→45.00 |
| | 4B | PT: Omitted conflict (State 1 v State 2) | 0.87→1.31; 3.01→4.14; 5.21→8.10; 8.67→9.36; 9.55→10.37 |
| | 4C | BCI: Lever press, conflict, State 1 | -10.13→-9.69; -8.49→15.04 |
| | 4C | BCI: Lever press, conflict, State 2 | -10.38→-9.75; -8.62→-7.86; -6.17→6.04 |
| | 4C | PT: Lever press, conflict (State 1 v State 2) | n.s. |
| Phase 3 State 1 (Risk-taking) | 4D | BCI: Responded (cue), Conflict | 0.31→0.88; 2.14→2.70; 4.97→10.00 |
| | 4D | BCI: Responded (cue), rewDS | 0.31→1.19; 1.76→10.00 |
| | 4D | PT: Responded (cue) (Conflict v rewDS) | n.s. |
| | 4E | BCI: Omitted, Conflict | 10.24→20.11; 22.06→22.50; 23.50→24.76; 25.95→28.03; 29.10→45.00 |
| | 4E | BCI: Omitted, punDS | 0.37→0.87; 2.32→2.95; 4.39→45.00 |
| | 4E | PT: Omitted (Conflict v punDS) | 1.88→2.89; 4.65→10.24; 22.37→22.81 |
| | 4F | BCI: Lever press, Conflict | -10.51→-9.69; -8.49→15.04 |
| | 4F | BCI: Lever press, rewDS | -10.07→15.04 |
| | 4F | PT: Lever press (Conflict v rewDS) | n.s. |
| Phase 3 State 2 (Risk-averse) | 4G | BCI: Responded (cue), Conflict | 9.25→10.00 |
| | 4G | BCI: Responded (cue), rewDS | 0.31→1.07; 5.35→10.00 |
| | 4G | PT: Responded (cue) (Conflict v rewDS) | 6.92→9.25 |

| | | | |
|---|---|---|---|
| | 4H | BCI: Omitted, Conflict | 0.31→1.00; 3.01→4.21; 5.27→6.28; 6.47→7.41; 7.54→8.48; 9.55→24.13; 24.45→45.00 |
| | 4H | BCI: Omitted, punDS | 0.37→0.94; 4.77→5.59; 9.11→32.87; 33.06→45.00 |
| | 4H | PT: Omitted (Conflict v punDS) | n.s. |
| | 4I | BCI: Lever press, Conflict | -10.38→-9.75; -8.56→-7.86; -6.61→-5.41; -5.28→6.04; 6.29→6.92 |
| | 4I | BCI: Lever press, punDS | -11.51→5.41; 5.54→8.18; 8.30→9.44; 9.56→12.27 |
| | 4I | PT: Lever press (Conflict v punDS) | n.s. |

**Table 2. Time periods (in seconds) where the photometry statistical tests are significant in Figure 4.** *Time 0 refers to cue onset for responded (A, D, G) and omitted (B, E, H) trials, and to the lever press for lever-press traces (C, F, I). On omitted trials the lever was inserted at 10 s and retracted at 40 s. State 1, Risk-taking; State 2, Risk-averse. BCI, Bootstrapped Confidence Intervals; PT = Permutation Tests.*

| Experimental Phase | Figure | Statistical Comparison | Significance time window (Seconds; 0 = Lever press) |
|---|---|---|---|
| Alcohol tests, State 1 (Risk-taking) | 5C | BCI: rewDS press, Saline | -6.22→12.68; 12.86→14.31 |
| | 5C | BCI: rewDS press, Alcohol 1.0 g/kg | -5.85→5.27 |
| | 5C | PT: rewDS press (Saline v Alcohol 1.0 g/kg) | 6.15→11.17; 11.55→12.49 |
| | 5D | BCI: Conflict press, reward, Saline | -1.45→4.20 |
| | 5D | BCI: Conflict press, reward, Alcohol 1.0 g/kg | -1.89→4.01 |
| | 5D | PT: Conflict press, reward (Saline v Alcohol 1.0 g/kg) | n.s. |
| | 5E | BCI: Conflict press, shock, Saline | -1.33→2.88; 4.58→5.39; 6.96→7.78; 8.09→8.72; 8.91→11.11 |
| | 5E | BCI: Conflict press, shock, Alcohol 1.0 g/kg | -2.33→-1.89; -1.58→3.13; 3.32→3.82 |
| | 5E | PT: Conflict press, shock (Saline v Alcohol 1.0 g/kg) | n.s. |

**Table 3. Time periods (in seconds) where the photometry statistical tests are significant in Figure 5.** *Time 0 refers to the lever press. All traces are from trials assigned to State 1 (Risk-taking), after the first 15 minutes of the test session. BCI, Bootstrapped Confidence Intervals; PT = Permutation Tests.*

| Experimental Phase | Figure | Statistical Comparison | Significance time window (Seconds; 0 = Cues turned on / Lever press) |
|---|---|---|---|
| Exp. 1 v 2 Phase 3 Responded (cue) | 6C | BCI: rewDS, Operant | 0.31→0.94; 2.08→10.00 |
| | 6C | BCI: rewDS, Pavlovian | 0.63→1.13 |
| | 6C | PT: rewDS (Operant v Pavlovian) | 1.82→10.00 |
| | 6C | BCI: punDS, Operant | n.s. |
| | 6C | BCI: punDS, Pavlovian | n.s. |
| | 6C | PT: punDS (Operant v Pavlovian) | n.s. |
| | 6C | BCI: Conflict, Operant | 0.38→0.82; 2.26→2.89; 8.11→10.00 |
| | 6C | BCI: Conflict, Pavlovian | 0.44→1.13 |
| | 6C | PT: Conflict (Operant v Pavlovian) | n.s. |
| Exp. 1 v 2 Phase 3 Outcome | 6D | BCI: rewDS, reward, Operant | -8.61→15.00 |
| | 6D | BCI: rewDS, reward, Pavlovian | -0.32→2.82 |
| | 6D | PT: rewDS, reward (Operant v Pavlovian) | -8.05→0.31 |
| | 6D | BCI: punDS, shock, Operant | -1.08→11.42 |
| | 6D | BCI: punDS, shock, Pavlovian | -0.26→11.48 |
| | 6D | PT: punDS, shock (Operant v Pavlovian) | -0.20→0.31 |
| | 6D | BCI: Conflict, reward, Operant | -2.71→15.00 |
| | 6D | BCI: Conflict, reward, Pavlovian | -2.33→-1.64; -1.45→2.82 |
| | 6D | PT: Conflict, reward (Operant v Pavlovian) | -0.20→0.31 |
| | 6D | BCI: Conflict, shock, Operant | -4.34→13.37 |
| | 6D | BCI: Conflict, shock, Pavlovian | -0.51→14.81 |
| | 6D | PT: Conflict, shock (Operant v Pavlovian) | -0.45→0.31 |
| Exp. 1 v 2 Phase 3 Omitted | 6E | BCI: rewDS, Operant | 10.24→32.55; 38.46→44.56 |
| | 6E | BCI: rewDS, Pavlovian | 10.43→11.25 |
| | 6E | PT: rewDS (Operant v Pavlovian) | 30.10→31.11; 35.26→40.35 |
| | 6E | BCI: punDS, Operant | 0.25→45.00 |
| | 6E | BCI: punDS, Pavlovian | 10.37→13.70; 40.35→45.00 |
| | 6E | PT: punDS (Operant v Pavlovian) | 0.18→0.69; 3.58→10.37; 17.41→40.35; 42.42→45.00 |
| | 6E | BCI: Conflict, Operant | 10.18→45.00 |

| | | | |
|---|---|---|---|
| | 6E | BCI: Conflict, Pavlovian | 10.43→13.57; 40.35→45.00 |
| | 6E | PT: Conflict (Operant v Pavlovian) | 17.59→21.93; 24.07→24.51; 29.22→40.29 |

**Table 4. Time periods (in seconds) where the photometry statistical tests are significant in Figure 6.** *Time 0 refers to cue onset for responded (C) and omitted (E) trials, and to the lever press (Operant) or outcome delivery (Pavlovian) for outcome traces (D). On omitted trials the lever was inserted at 10 s and retracted at 40 s. BCI, Bootstrapped Confidence Intervals; PT = Permutation Tests.*

| Experimental Phase | Figure | Statistical Comparison | Significance time window (Seconds; 0 = Cues turned on / Lever press) |
|---|---|---|---|
| Exp. 1 v 2 Phase 2 Responded (cue) | S6B | BCI: rewDS, Operant | 0.38→0.88; 1.76→10.00 |
| | S6B | BCI: rewDS, Pavlovian | n.s. |
| | S6B | PT: rewDS (Operant v Pavlovian) | 2.39→10.00 |
| | S6B | BCI: punDS, Operant | 6.23→6.79; 7.80→9.37 |
| | S6B | BCI: punDS, Pavlovian | n.s. |
| | S6B | PT: punDS (Operant v Pavlovian) | 6.23→10.00 |
| Exp. 1 v 2 Phase 2 Outcome | S6C | BCI: rewDS, reward, Operant | -10.60→15.00 |
| | S6C | BCI: rewDS, reward, Pavlovian | -0.75→1.95; 8.97→15.00 |
| | S6C | PT: rewDS, reward (Operant v Pavlovian) | -8.03→7.59; -6.72→0.38 |
| | S6C | BCI: punDS, shock, Operant | -6.65→5.85 |
| | S6C | BCI: punDS, shock, Pavlovian | 0.13→12.74 |
| | S6C | PT: punDS, shock (Operant v Pavlovian) | -4.33→0.31 |
| Exp. 1 v 2 Phase 2 Omitted | S6D | BCI: rewDS, Operant | n.s. |
| | S6D | BCI: rewDS, Pavlovian | 10.55→12.75; 40.47→41.17 |
| | S6D | PT: rewDS (Operant v Pavlovian) | n.s. |
| | S6D | BCI: punDS, Operant | 0.37→0.87; 10.18→45.00 |
| | S6D | BCI: punDS, Pavlovian | 10.24→15.77; 20.49→24.32; 26.65→45.00 |
| | S6D | PT: punDS (Operant v Pavlovian) | -1.45→-0.89; 15.65→27.34; 27.84→29.29; 29.85→30.35; 33.94→34.38; 36.39→37.52; 37.71→38.78 |

**Supplementary Table S1. Time periods (in seconds) where the photometry statistical tests are significant in Supplementary Figure 6.** *Time 0 refers to cue onset for responded (B) and omitted (D) trials, and to the lever press (Operant) or outcome delivery (Pavlovian) for outcome traces (C). On omitted trials the lever was inserted at 10 s and retracted at 40 s. BCI, Bootstrapped Confidence Intervals; PT = Permutation Tests.*
